# Time-Aligned Hourglass Gastrulation Models in Rabbit and Mouse

**DOI:** 10.1101/2022.11.13.516304

**Authors:** Y Mayshar, O Raz, S Cheng, R Ben-Yair, R Hadas, N Reines, M Mittnenzweig, O Ben-Kiki, A Lifshitz, A Tanay, Y Stelzer

## Abstract

The hourglass model describes the convergence of species within the same phylum to a similar body plan during development, yet the molecular mechanisms underlying this phenomenon in mammals remain poorly described. Here, we compare rabbit and mouse time-resolved differentiation trajectories to revisit this model at single cell resolution. We modeled gastrulation dynamics using hundreds of embryos sampled between gestation days 6.0-8.5, and compare the species using a new framework for time-resolved single-cell differentiation-flows analysis. We find convergence toward similar cell state compositions at E7.5, underlied by quantitatively conserved expression of 76 transcription factors, despite divergence in surrounding trophoblast and hypoblast signaling. However, we observed noticeable changes in specification timing of some lineages, and divergence of primordial germ cells programs, which in the rabbit do not activate mesoderm genes. Comparative analysis of temporal differentiation models provides a new basis for studying the evolution of gastrulation dynamics across mammals.

## INTRODUCTION

Early mammalian development follows a generally conserved sequence of events resulting in extraordinarily similar embryos at the time of organogenesis, representing a so-called evolutionary hourglass effect^1-3^. The process of gastrulation involves the formation of the embryonic germ layers from the pluripotent epiblast, and laying out of the basic embryonic axes. This critical stage of development has been mainly characterized in the mouse model, in which the developing blastocyst takes on the form of a cup shape, or egg cylinder. The mouse gastrula is however, highly distinct among vertebrates, as most mammals begin gastrulation as a planar embryonic disc. Such gross structural disparity is expected to have dramatic effects on gastrulation by shaping cellular mechanics and spatiotemporal interactions. Moreover, wide variation has been observed between species in relation to implantation strategies, and the development and orientation of the extraembryonic tissues^4,5^. The rabbit stands out among possible alternative mammalian model organisms by presenting many of the advantages of the mouse, namely, relatively short gestation, and large litters that can be accurately timed. Importantly, the rabbit was shown to more closely resemble human development, in particular with respect to the specification of primordial germ cells (PGCs)^6,7^. Interestingly, implantation of rabbit embryos is preceded by a period of rapid growth as an expanded blastocyst^8,9^, and in this form gastrulation is initiated^10^. Further, while anti-mesometrial implantation occurs at embryonic day (E)7, placental (mesometrial) implantation occurs only at E8^11^, close to the time of somitogenesis. This is in contrast to the mouse, in which implantation occurs early after hatching of the blastocyst at around E4.5, long before the onset of gastrulation.

Single cell transcriptomics and multi-omics are recently being used to map transcriptional and chromatin states in early embryonic development^7,12-22^. This resulted in comprehensive atlases that greatly enrich and refine previous imaging-based data by characterizing precisely transcriptional programs at high cellular resolution. Merging inferred cell states into a manifold model facilitates the inference of cellular differentiation dynamics, at first using computational tools searching for parsimonious differentiation trajectories^23,24^. But the highly parallel and complex nature of embryonic development, and gastrulation in particular, calls for refined strategies to map precisely multi-lineage dynamics of the ensemble of single cells constituting the developing embryo. We recently developed such an approach by sampling single cells from individual embryos, and by inferring a differentiation flow model that describes how single cells coordinately change their manifold state over time^14^. This approach can also be instrumental to elucidate the intrinsic and extrinsic function of genes in embryonic cell specification^25^. Together with technologies for mapping cell lineage trees (Reviewed in^26^), and spatial transcriptomics^27-29^, this can be even further refined to obtain a true holistic view of embryonic development.

Importantly, embryonic atlases were recently described from precious and limited primate^30,31^ and human^20,32^ samples, but the temporal resolution in these systems is limited. Here, we adapt the rabbit as a second high resolution single cell model for mammalian gastrulation. We generated a time resolved single-embryo, single-cell model from over 100 embryos, and used it for comprehensive synthesis of morphological and molecular classification of rabbit embryonic stages. We define the manifold alignment problem as the evolutionary task of matching molecular states (e.g. cell types) and differentiation dynamics between related species, and describe how to approach this problem when comparing mouse and rabbit models. This leads to characterization of striking resemblance in gastrulation programs between the two species, but also uncovers key differences in lineage coordination and differentiation timing, setting the stage for understanding how conservation of the mammalian intrinsic cell state repertoire can be compatible with changes in morphology and extrinsic signaling.

## RESULTS

### Charting the rabbit single cell gastrulation manifold using 120 individually profiled E6.0– E8.5 embryos

While overall gestation of the rabbit is significantly longer than that of the mouse (∼30 vs ∼19 days), major early gestational milestones such as somitogenesis, limb, and eye development occur at highly similar times^33^. Also, both species begin gastrulation shortly after E6.0. However, despite the similarity in timing of gastrulation, and in form during organogenesis, several key differences in the process of gastrulation exist between the species: (i) The rabbit embryo is much larger than the mouse during all corresponding developmental stages. (ii) Uterine implantation, which in the mouse occurs prior to gastrulation, only occurs gradually in the rabbit and at much later stages between embryonic days E7-8. (iii) The egg cylinder shape of the mouse facilities asymmetric interaction between the epiblast and extraembryonic tissues (distal-proximal axis), as opposed to the disc shape of the rabbit embryo which in practice lacks such an axis. (iv) Establishment of the mouse anterior-posterior axis involves coordinated signals from the anterior visceral endoderm (AVE, corresponding to rabbit’s anterior hypoblast) and the extraembryonic ectoderm (ExEc). However, in the rabbit, the part of the ExEc that is in contact with the embryo proper (polar trophectoderm, also known as Rauber’s layer) deteriorates and disappears rapidly at the onset of gastrulation^34^ and is therefore unlikely to play a major signaling role at this stage.

To better understand the conservation of the gastrulation program and its archetypical outcome given such a diverse context, we sought to generate a time-resolved atlas of rabbit gastrulation using a similar approach we previously implemented in the mouse^14^. To this end, we performed scRNA-seq^35^ on 160K cells of 120 individual embryos including their immediate surrounding extraembryonic tissue. Each embryo was imaged and assigned a morphological stage^36^, and profiles of single cells were associated with their corresponding embryo. Litters were timed to ensure adequate representation of the structural diversity from pre-gastrulation up to the 12 somite stage (**Figures 1A, S1, S2A-D**). Following quality control, we retained 130K profiles of sufficient coverage (**Figure S2A**, median UMI count of 7,142) and constructed a transcriptional manifold model using the metacells2 algorithm^37^. We have extended the rabbit gene annotation to the yet unannotated oryCun3.0 genome assembly using the previous state-of-the-art oryCun2.0 gene annotation as well as the mouse mm9 and Human hg19 gene annotations. Furthermore, we account for any unannotated transcriptional activity that can be mapped to oryCun3.0, adding 120K putative 3’-end loci to the OryCun3.0 25K annotated TSSs (**Figure S2G**). In total we defined 3,584 metacells (MCs), facilitating precise quantitative and comparative analysis of the represented transcriptional states (**Figure 1B**). We annotated ExEc and extraembryonic endoderm (ExEnd) cells and analyzed them separately as elaborated below. We then annotated the embryo manifold using 41 different cell types/states. The entire model is accessible interactively at tanaylab.weizmann.ac.il/rabemb_wex. For subsequent analysis of embryo temporal dynamics, embryos were excluded for either insufficient embryonic cells (<20 cells, n=6), or obvious outliers in cell type composition (indicating a technical error, e.g. embryos with only posterior cell types, n=6).

**Figure 1.**
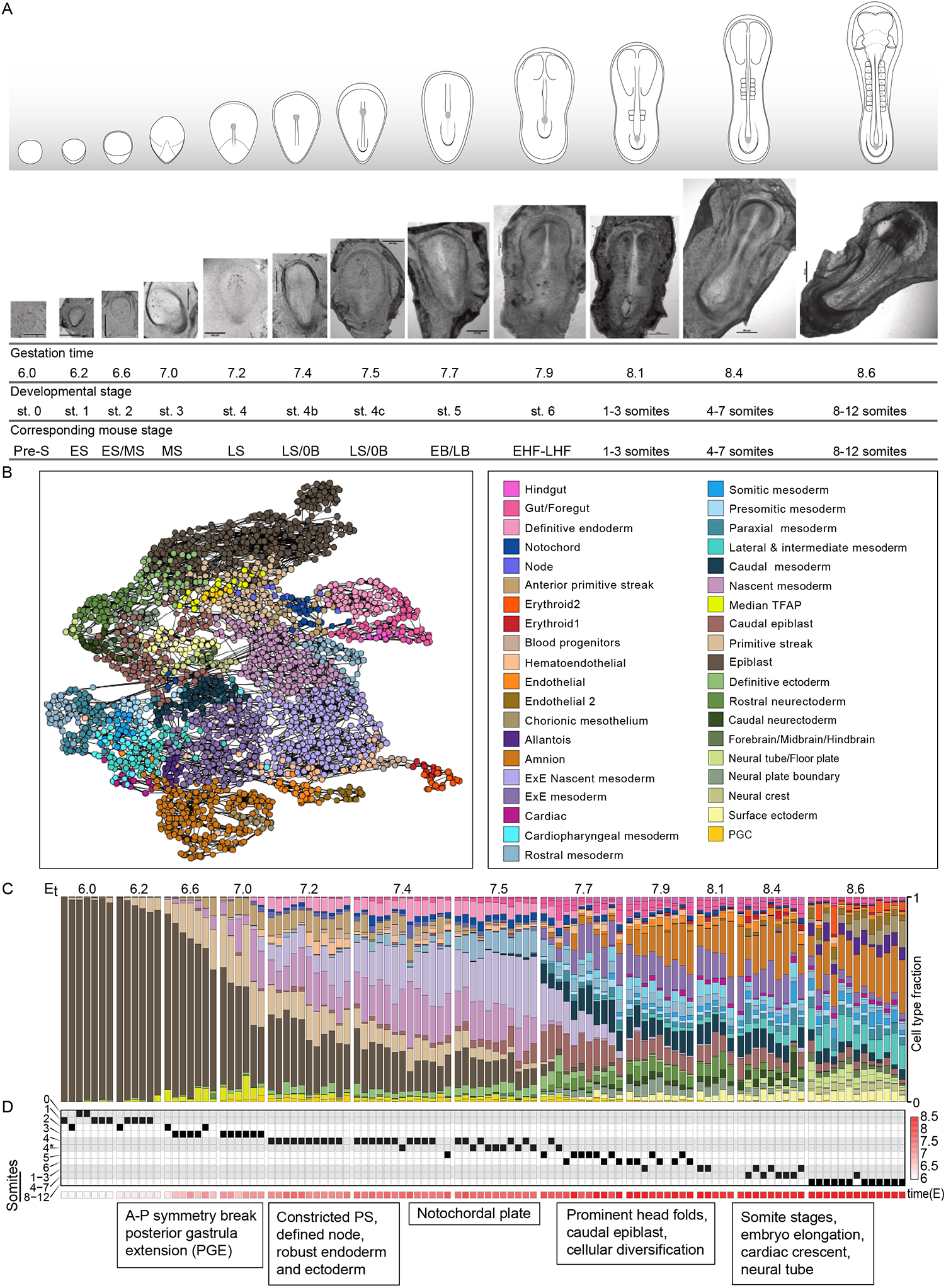
Generation of a temporal rabbit single cell transcription manifold. (A) Morphological characterization of rabbit embryos analyzed in this study. Shown is a schematic representation of the main morphological features observed in each embryonic stage, alongside an image of a representative embryo. Gestation time reflects a single idealized time for each embryo time group. Images are uniformly scaled. Scale bar = 500um. (B) Color coded 2-D UMAP projection of the metacells (MCs) graph representing the entire transcriptional manifold. MCs (nodes) are connected by edges, indicating the most similar neighbors for each MC. (C) Cell type distribution per embryo, arranged according to intrinsic transcriptional rank and separated into 12 characteristic age groups. (D) Morphological staging of individual embryos, indicated by a black square. Embryos that could not be assigned a stage (lacking proper image) are left empty. Color bar indicates the actual gestation time of each embryo, calculated from the time of mating. Some of the most distinctive morphological and cell state events are noted in the boxes below.

### Refining rabbit embryonic stages by integrating morphology and transcriptional analysis

To describe rabbit gastrulation on an absolute temporal axis, we performed morphology-based ranking of the embryos and ranking by k-NN similarities of single-cell profiles (**Figures S1 and S2H**). Both measurements showed high concordance and generally matched the respective litter’s absolute gestation time (**Figure S2E-F**). We then defined twelve temporal bins comprising groups of embryos with similar cell state compositions (see **Methods**), which were used in subsequent modeling and refinement of the current morphology-based nomenclature (**Figure 1C**). Under the assumption that all embryos represent discreet times along a shared process, each bin was manually assigned a representative time considering the absolute gestation time of embryos comprising it, defined as E_t_ (See **Methods**, and **Table S1**). Each embryo was also assigned to a specific morphological developmental stage, resulting in a multi-tiered annotation of the rabbit gastrulation continuum (**Figure 1D**). We find that cell type composition changes in many cases reflect visible morphological events, such as; a) the elongation of the embryo from its initial circular form that coincides with the appearance of the mesoderm and anterior primitive streak; b) the appearance of a narrowed primitive streak structure (E_t_7.2, stage 4) together with definitive endoderm and ectoderm, hematoendothelial cells, and expansion of the extraembryonic nascent mesoderm, and, c) the brief window of time between the first observation of the notochordal process anterior to the node (E_t_ 7.5, stage 4c), and its full anterior extension (E_t_7.7, stage 5), that is accompanied by an explosion of advanced cell types and disappearance of the nascent mesoderm. Cell type composition analysis also allows direct comparison with similar mouse analysis, providing an empirical measure of corresponding developmental stages across species. In summary, single cell analysis of single rabbit embryos provided us with high resolution description of the rabbit gastrulation stages. It thereby set the stage for quantitative modeling of the differentiation process, transforming the pluripotent epiblast into the rich repertoire of embryonic lineages during 2.5 days from E6.0 to E8.6.

### A network flow model tracks rabbit gastrulation dynamics in absolute time

To infer a differentiation model from the single-cell and single embryo rabbit dataset, we used an improved version of our network flows algorithm that was initially demonstrated in the mouse^14^ (see **Methods**). The algorithm resolved differentiation flows for metacells distributed over the twelve time-bins, balancing similarities between expression states and estimation of cell proliferation rates (**Figures 2, S3A, and S3B**). The latter was performed by computing co-expression of S-phase and M-phase related genes, and quantifying a distinctive non-proliferating cell subpopulation (**Methods, Figure S3C**). Consistent with the extremely high proliferation rate in gastrulating rabbit embryos (estimated doubling every 6-7 hours^9^) we observe that almost all cells express the cell cycle programs. Nevertheless, some metacells of the endoderm lineage, notochord, and cardiomyocytes were linked with considerably lower expression of S- and M-phase related genes, suggesting a slow-down in proliferation rates in these cell states (**Figure S3D**). Estimations of the fraction of non-dividing cells per cell state and time were used to calibrate differential growth rates and growth loss estimation for the flows ‘mass conservation’ constraint (See **Methods**). Yet, for some extraembryonic lineages, we could not assume such conservation due to incomplete or biased sampling. Namely, the rabbit choriovitelline placenta tissue is extensive and lacks visible borders by standard light microscopy. Therefore, rabbit embryo resection entails arbitrary removal of much of the yolk sac endoderm and related extraembryonic mesoderm, which extends radially from the embryo and thus progressively undersampled. Hence, analysis of cellular hierarchy was excluded for hematopoietic, ExEnd, and trophectoderm lineages (even though the timing of their sequential appearance remains highly informative, as discussed below). Apart from these lineages, the rabbit flow model uncovers the exceptionally coordinated process of differentiation in the ectoderm, mesoderm, and endoderm with high temporal and quantitative precision. Importantly, this model is based on similar assumptions to those used to create the analogous mouse model and therefore allows direct quantitative alignment between the two species.

**Figure 2.**
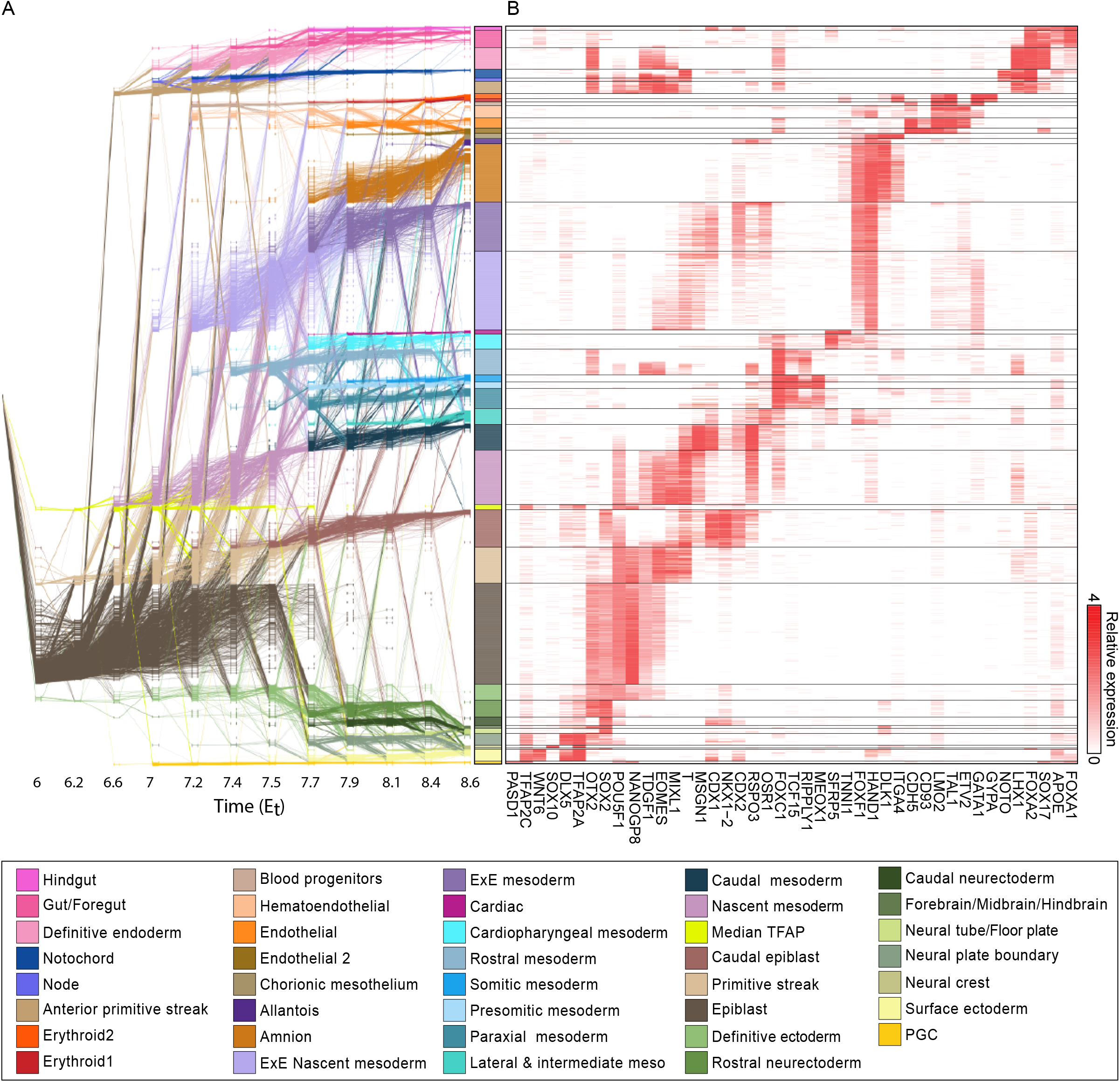
Rabbit gastrulation network flow model. (A) The gastrulation network flow model consists of MCs (nodes in rows) distributed in time (x-axis), and flows (edges) that link MCs between adjacent time points. The first time-point represents a common source for all MCs. Annotation (color coded) relies on marker expression and flow-based fate mapping. (B) Heatmap showing relative expression (log2) of key lineage-specific genes. Extraembryonic endoderm and ectoderm lineages are presented separately, see Figure 4.

### Manifold alignment uncovers highly conserved gastrulation states in rabbit and mouse

Given fully time resolved manifold and flow models for rabbit and mouse gastrulation, we wished to define a framework for their principled comparison. Previous comparative single cell atlas work analyzed either highly diverged species with non-alignable genomes^38,39^, or in the case of mammals, using lower resolution data^19,30,40,41^. These analyses therefore were restricted to qualitative and coarse grain comparisons. To facilitate a high-resolution comparative analysis, we developed a manifold alignment algorithm that simultaneously computes matching states between mouse and rabbit while adjusting functional gene orthology. Our approach begins with a naïve assumption of complete functional conservation of 9,748 orthologous gene pairs for which significant transcription is detected in both species. This pairing was used for initial approximation of the similarity between all metacells in the two manifolds (**Figure S4A**), identifying 79 reciprocally best orthologous metacell (RBOM) pairs (**Figure S4B**). Notably, the identification of such states is a highly non-trivial result, showing that the two manifolds are indeed alignable over a very rich collection of transcriptional states. Using the 79 RBOMs as a new common reference, we next recomputed gene pairing between the species. This involved; a) identifying sequence orthologs that are transcriptionally diverged (**Figure S4C**); b) adding constant fold factor correction for orthologs that show species-specific constant bias (**Figure S4D**); c) discovering new putative ortholog associations for un-annotated (**Figure S4E**), or ambiguously aligned (**Figure S4F**) rabbit genes that are linked uniquely to mouse genes. Following functional orthology adjustments, we recomputed rabbit-mouse similarity over all manifold states. This procedure uncovered a highly specific positive correlation between related cell type pairs and anti-correlation for unrelated pairs (**Figure 3A**). We note that while overall alignment between the two manifolds is specific and inclusive, it can be of variable precision with significant gene divergence observed even for highly correlated states. Notably, the matching of PGC states between the species was lower than observed for other states (**Figure 3B**). Manifold state alignment can also represent broad similarities between related groups of states rather than a one-to-one state correspondence. For example, the rabbit ‘median TFAP’ state is not directly matched by a corresponding mouse annotation, but still shows some similarity to the mouse epiblast, definitive ectoderm, primitive streak, and PGCs (**Figure 3A**).

**Figure 3.**
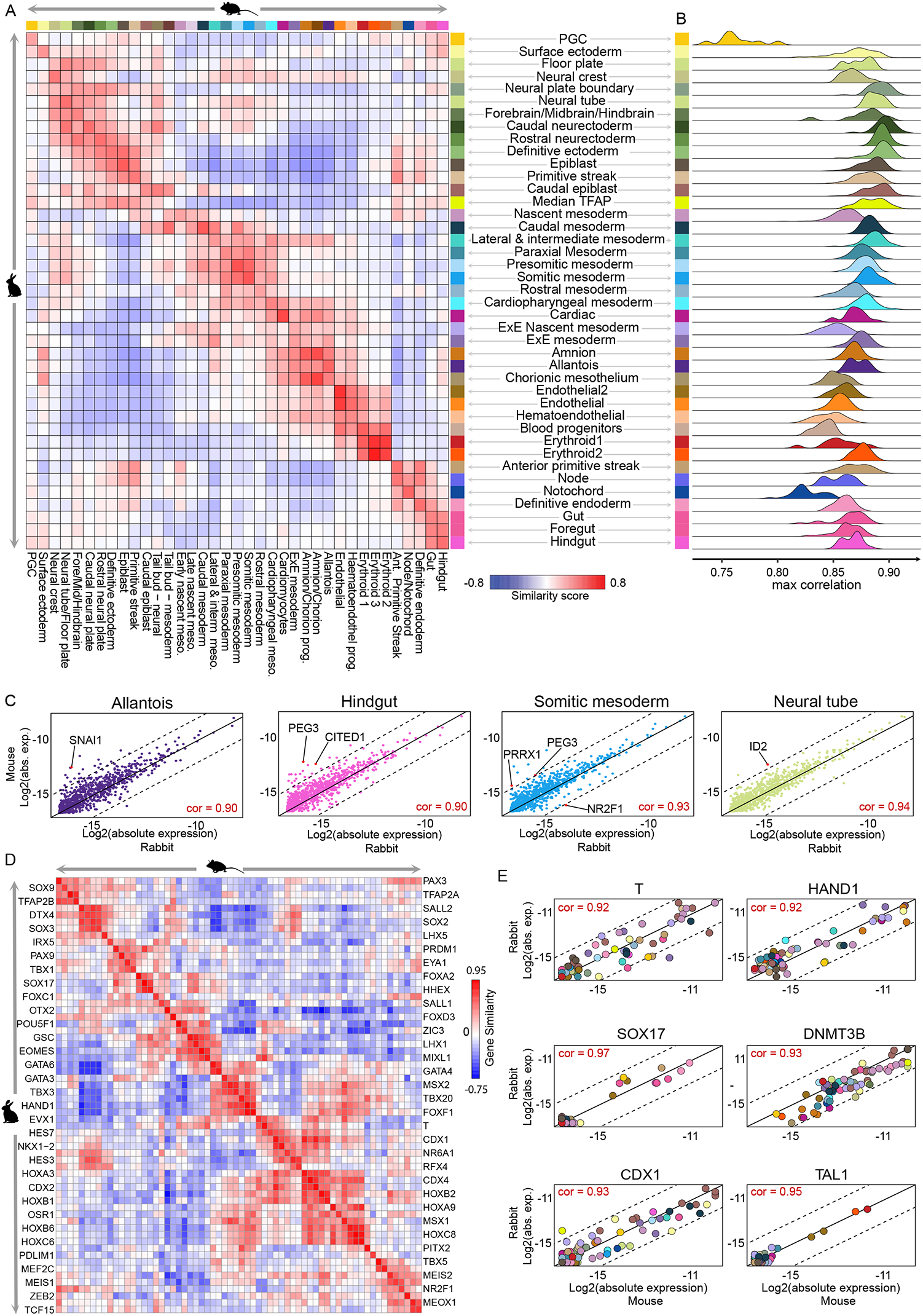
Rabbit/Mouse cross-species manifold alignment. (A) Comparison of independently annotated rabbit and mouse transcriptional states according to similarity of global gene expression profiles (here, based on 955 genes that are the intersection of each manifolds set of feature genes). (B) Accumulated maximal correlation between rabbit MCs of each type, to any MC in the mouse manifold. (C) Comparable gene expression in selected cell types. Each point indicates absolute expression levels in rabbit and mouse for each gene (log2 of UMI frequency, x- and y-axes, respectively, averaged by cell type). (D) Gene-gene correlations between reciprocally best orthologous metacells (RBOMs). Shown are most highly correlated transcription factors between species. (E) MC-MC correlations between RBOMs for a number of representative genes (following linear correction, see Methods).

### A conserved gene regulatory core of transcription factors underlies gastrulation

Aligned manifold models at metacell resolution facilitate a fully quantitative comparison of the gastrulation transcriptional states. As exemplified in **Figure 3C**, the expression of gastrulation genes is, in most cases, within two-fold difference between the aligned species states. This analysis also allows the detection of genes that escape such conservation, such as SNAI1 for mouse allantois, NR2F1 for rabbit somitic mesoderm, and more (**Figure 3C** and **Table S3**). Next, we sought to systematically characterize the core gene regulatory conservation between the two gastrulation manifolds. We computed a cross-species gene by gene correlation matrix and searched for pairs of genes (not necessarily sequence orthologs) that were reciprocally best correlated. Of these, 147 gene pairs were found to be best reciprocal sequence orthologs, the majority of which (n=76) code for transcription factors (TFs) (**Figures 3D-E**. See complete list in **Table S2**). These TFs constitute a massive core of conserved regulatory logic that drives gastrulation. This logic is rapidly emerging over time, where 72% of the core TFs are significantly expressed (log2(UMI frequency)>-14) in at least one metacell by E_t_7.5. In almost all cases, the identity of these early and concurrently induced TFs is still unmatched with specialized downstream programs (i.e. characterized by structural and functional genes). The deep conservation of the gastrulation TF core is particularly surprising given the high degree of regulatory redundancy in the system, with multiple TFs active and regulating each lineage^14^. This points toward a finely-tuned quantitative balance that goes beyond redundancy, suggesting such balance is crucial for coordinating the complex cellular multifurcation that installs the blueprint of cellular lineages during gastrulation.

### Partial convergence of rabbit and mouse extraembryonic endoderm programs

The conservation of gastrulation transcriptional states is compatible with the known morphological convergence of early mammalian embryos termed the phylotypic stage, and the related hourglass model. In contrast to this conservation, the rabbit and mouse extraembryonic tissues highly diverge in their structure, with the rabbit displaying typical planar mammalian hypoblast and trophoblast, largely distinct from the mouse (**Figure 4A**). To understand how such morphological divergence may potentially impact signaling between extraembryonic and embryonic tissues and hence their role in axial patterning^42^, we created a rabbit ExEnd manifold (**Figure 4B**) and compared its states to the mouse counterpart (**Figures 4C-D**). This identified a program characterized by expression of Laminin (e.g. LAMA1, LAMB1), which characterized mouse parietal endoderm (PE), and in the rabbit included both PE, and hypoblast (Hyp) states (the latter showing higher expression of the pluripotent factor ESRRB). A second matching behavior was defined by strong expression of Apolipoproteins (APO-A/B/C), involving rabbit yolk sac endoderm (YSE) states^43^, and in mouse embryo-associated visceral endoderm (VE) and extraembryonic VE. We also detected a subpopulation of rabbit hypoblast cells expressing the TFs EOMES, LHX1, HHEX, and the BMP antagonist CER1, together with relatively lower WNT3. This subpopulation therefore exhibits a transcriptional repertoire closely reminiscent of the mouse anterior visceral endoderm (AVE), and was termed anterior hypoblast^44,45^. As all rabbit epiblast cells are initially in direct contact with the hypoblast cell layer, specialized hypoblast programs were previously suggested to underlie the anterior-posterior polarization of the embryo^46^. Experimentally, by resection of the rabbit hypoblast, it was demonstrated to play an inhibitory role, confining the domain of the primitive streak to its stereotypic form^47^. Importantly, the convergence toward common functional programs in the endoderm of both species occurs even though some of the regulatory machinery is diverged, as is the case for EPAS1 and MYC, two key TFs that follow reversed patterns (**Figure 4D**).

**Figure 4.**
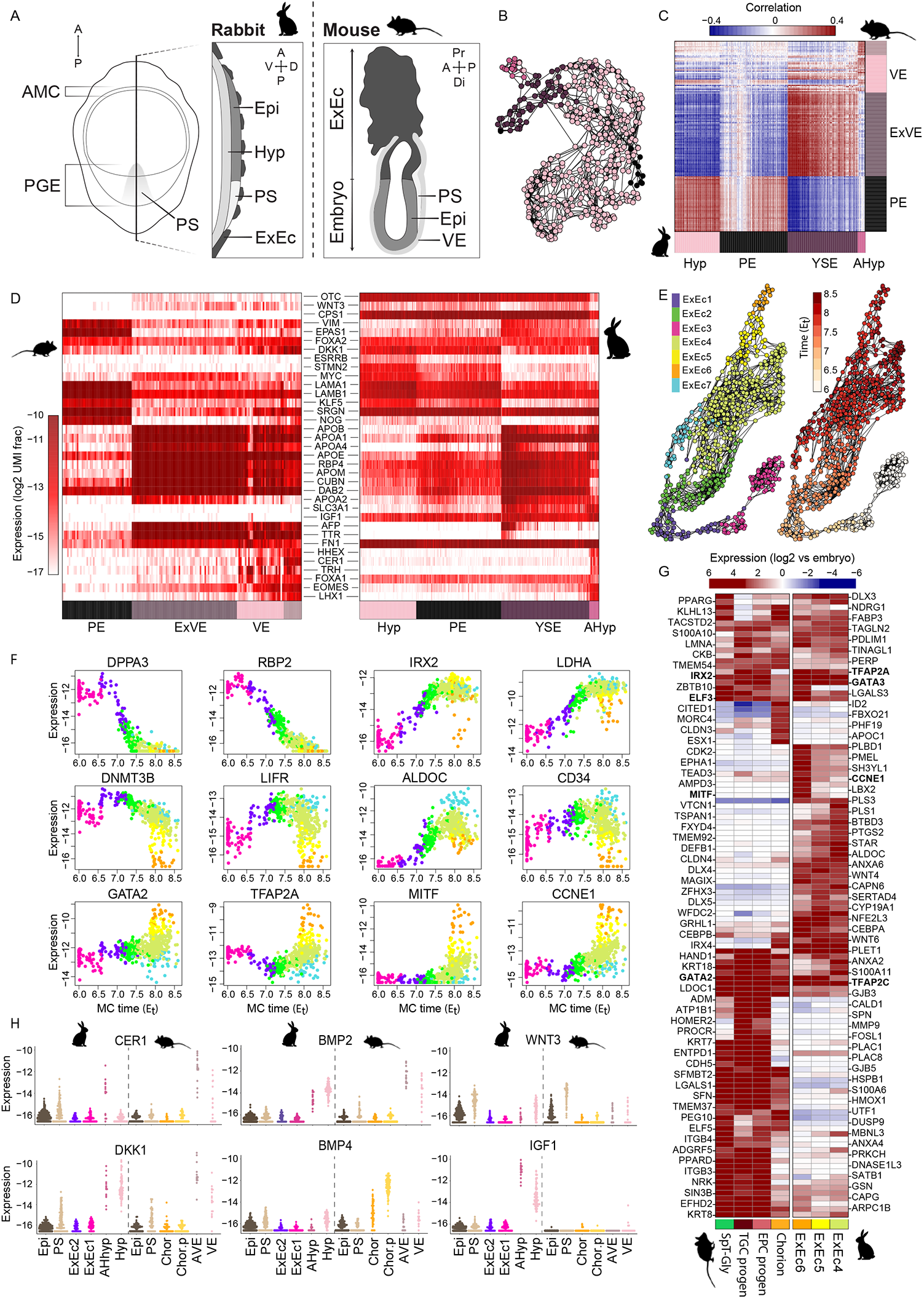
Similarities and differences in extraembryonic endoderm and ectoderm. (A) Schematic of structural organization of late streak rabbit and mouse embryos. Midsagittal sections of mouse and rabbit are shown alongside an en-face view of the rabbit embryo. The planar form of the hypoblast and trophoblast observed in rabbits diverges from the cup-shaped mouse embryo. (B, E) Color coded 2-D UMAP projection of the MC graph representing the extraembryonic endoderm and ectoderm manifolds, respectively. MCs (nodes) are connected by edges, indicating the most similar neighbors for each MC. (C) MC-MC correlation matrix of rabbit and mouse extraembryonic endoderm. (D) Side by side expression heat map (log2 of UMI frequency) of highly variable genes in rabbit and mouse extraembryonic endoderm. (F) Temporal expression of key genes during ExEc development. Each dot represents a single MC. (G) Heatmap showing relative expression of most highly variable genes within the ExEc (log2 fold-change relative to the embryonic average). (H) Expression of key signaling factors during early gastrulation. Each dot represents a single MC. Only MCs with calculated time of E6-6.5 are shown. AMC – anterior marginal crescent; PGE – posterior gastrula extension; PS – primitive streak; Epi – epiblast; Hyp – hypoblast; ExEc – extraembryonic ectoderm; VE – visceral endoderm; AVE – anterior visceral endoderm; ExVE – extraembryonic visceral endoderm; YSE – yolk sac endoderm; AHyp – anterior hypoblast; PE – parietal endoderm; Chor – chorion; Chor.p – chorion progenitors.

### Delayed and diverged rabbit trophoblast differentiation

In contrast to the overall transcriptional convergence of ExEnd states, the rabbit extraembryonic ectoderm (ExEc) states (**Figures 4E**) show little resemblance to those of the mouse (**Figures S5A-B**). The early bifurcating and diverging mouse chorion and ectoplacental cone (EPC) populations^48,49^ could not be observed in rabbit ExEc. Instead, we found highly homogeneous states that gradually reduced DPPA3, RBP2 and increase IRX2, LDHA, among many other genes through E_t_7.5 and until E_t_8.0 (stages 4b-6) (**Figures 4F** - top, and **S5C-D**). Only following E_t_8.0 (somite stages), coinciding with time of mesometrial implantation^11^, diversification of the ExEc programs become evident, along a gradient spanned by diversifying expression of genes including GATA2, TFAP2A, MITF, and CCNE1 (**Figure 4F** - bottom) and inverse regulation of DNMT3B, LIFR, ALDOC, and CD34 (**Figure 4F** - middle). Comparison of this spectrum of late rabbit ExEc states to the mouse chorion and EPC states (**Figure 4G**) reveals important common features, including several TFs (TFAP2C, IRX2, ELF3, GATA3) and structural genes such as PLET1 and LMNA. Nevertheless, the differentiation gradient of late rabbit ExEc states did not associate strongly with either mouse chorion or derivatives of the ectoplacental cone lineage. This may either suggest dramatically altered placental transcriptional states, or that rabbit differentiation toward functional placenta precursors initiates beyond the 8-12 somite stage (E_t_8.6).

### Differential expression of growth factors in early extraembryonic tissues

In the mouse, reciprocal interactions between the epiblast and ExEc that involve BMP4, NODAL, and WNT3, regulate the induction of the primitive streak (PS)^50-52^. Similarly, inhibitory signals produced by the mouse AVE, namely, the BMP and WNT inhibitors CER1 and DKK1, are critical for proper anterior-posterior axis formation^45^. Analysis of the rabbit hypoblast and ExEc states at the relevant time, prior to E_t_6.5, showed conservation of BMP2 and WNT3 expression in the hypoblast. Notably, while CER1 is enriched in the anterior hypoblast, DKK1 appears to be expressed throughout the hypoblast (**Figure 4H**). BMP4 however, is not expressed in early ExEc cells, and only at low levels in the hypoblast. This, despite its robust expression in the embryo proper at later time stages, suggests the predominance of the more robustly expressed BMP2 for early patterning (**Figures S5A, F**). Finally, we find IGF1 to be highly expressed in the rabbit hypoblast, while in the mouse IGF1 expression is confined to the much later endothelial cells (also shared in the rabbit) (**Figure S5G**). In line with this finding is the demonstrated role of IGF1 in activation of embryoblast specific signal transduction^53^, and maturation of ex utero cultured rabbit blastocysts^54^. Together, this analysis suggests a minimal signaling role for the rabbit ExEc in embryonic patterning (consistent with the deterioration of the Rauber’s layer), and supports previous findings regarding the prominent role of hypoblast polarity. The detailed temporal expression of signaling genes (**Table S3**) complements previous efforts using traditional molecular techniques^46,54-56^, and provides a basis for further understanding intercellular signaling developmental regulation in non-rodent gastrulating embryos.

### Aligned rabbit and mouse gastrulation time axis highlights an hourglass-like bottleneck

Following alignment of the rabbit and mouse gastrulation manifolds, we next sought a principled strategy for comparing the two differentiation *processes*, as represented by our models. We therefore tested whether the independently determined rabbit and mouse gastrulation “clocks” could be aligned over a common time axis. To eliminate potential cell type annotation bias we generated a unified representation of rabbit/mouse cell state distributions (See **Methods**), and in this manner computed cross-species embryo cell-state frequency similarities over time (**Figure 5A**). The resulting similarity matrix revealed a stereotypical structure in which pre-gastrulation states are aligned but not synchronized, leading toward a bottleneck at approximately E_t_7.5 in both species, followed by a more synchronized gastrulation process with potential gradual loss of coherence. This analysis showed that overall we can use the absolute time axes of the two species for comparing their gastrulation processes. It also clearly demonstrates divergence and compensation in some key stages, particularly highlighting the earlier and more gradual emergence of PS populations in the mouse (**Figures 5B, C**). Remarkably, although the rabbit PS emerges later, it then converges to embryonic frequencies that closely match those observed in mouse. We next performed detailed comparison of the conserved and diverged gene expression signatures in the epiblast and PS of both species (**Figure S6A, B**). This highlighted the conservation of most of the previously characterized PS TFs and signaling genes, but also detected intriguing divergence of several developmental genes such as GATA4, FGF4, and POU3F1. This data further demonstrates conservation in the establishment of the core PS program, despite sizing, contextual and morphological differences between rabbit and mouse pre-gastrulation embryos.

**Figure 5.**
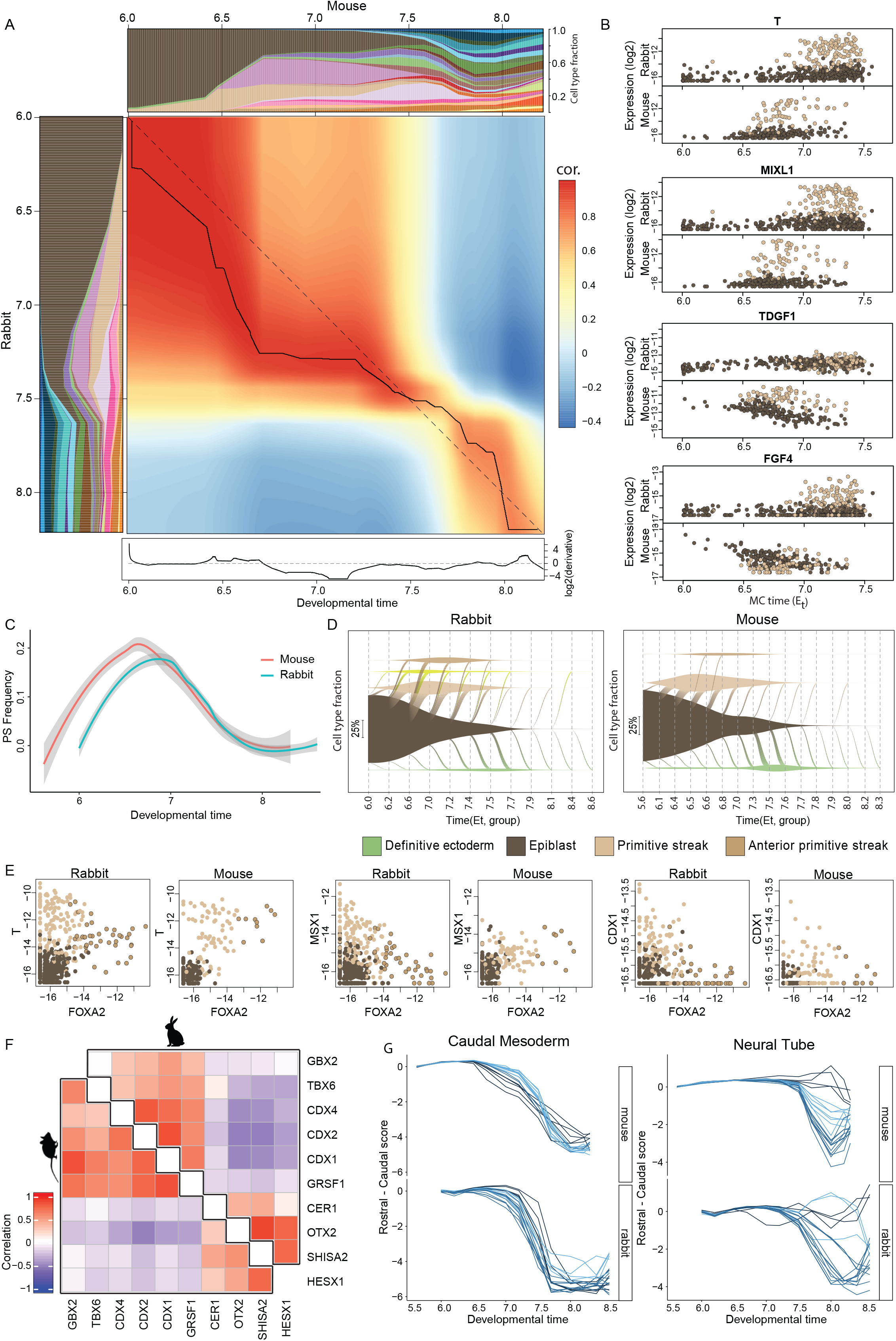
Interspecies temporal alignment. (A) Heatmap of interspecies correlation of cell type composition over time. For comparability, rabbit MC identities were derived de novo by the most similar mouse annotation according to gene expression (see Methods). Dynamic time warping (DTW) is plotted as a solid line along the diagonal (dashed), explaining differences as temporal compression/stretching. Plotted below, the DTW derivative, highlighting the temporal segments of major changes. (B) Temporal expression of key genes during epiblast to PS transition. Each dot represents a single MC. (C) Temporal alignment of PS frequency in mouse and rabbit. Lines represent Loess smoothing, standard error bounds indicated by shading. (D) Vein plots describing the continuous transition of cell types from the epiblast (represented by diagonal flows spanning time points), and the dynamic relative frequencies of these cell types in the embryo over time (vein width on the y-axis). Vein connection width represents flow flux at each time point. (E) Temporal expression of key genes in epiblast, PS and APS. Each dot represents a single MC. (F) Correlation in rabbit and mouse of, Rostral genes: CER1, OTX1, SHISA2, HESX1, and Caudal genes: GBX2, TBX6, CDX1,2,4, GRSF1. (G) Rostral-Caudal score of individual MC trajectories of terminal types, as indicated.

Interestingly, initiation of gastrulation between the species is also characterized by clear differences in the induction of the anterior primitive streak (APS). In contrast to the gradual acquisition of APS through an intermediate PS state in the mouse, rabbit APS rapidly accumulates and appears to be induced directly from the epiblast (**Figure 5D**). **Figure 5E** shows that mouse PS states co-expressing T/Brachyuri (key TF of the PS) and FOXA2 (early endoderm TF) are not matched in the rabbit manifold. Instead, rabbit epiblast differentiation toward the PS quickly induces MSX1 (mesodermal TF) and CDX1 (caudal homeobox), distinguishing itself from high FOXA2 states linked with the APS. Together, these results indicate that rabbit endoderm may be established without transiently activating a meso-endo like program. This also serves as an example for conserved states that can be developmentally reached, even though the trajectories leading to them have changed during evolution.

### Conserved rostral-caudal signatures in morphologically diverged gastrulas

Following the observation of species-specific timing of PS bifurcation, we next tested systematically whether this might further propagate to affect the later rostral-caudal program. To this end, we searched for conserved caudal and rostral gene regulatory modules (**Figure 5F**), and computed for each metacell a rostral/caudal score by calculating the expression ratio between the two. We then used the flow models to traceback the differentiation trajectories of metacells, grouped by type, and charted the rostral/caudal scores of each trajectory over time (**Figures 5G and S7A-H**). This showed remarkable conservation in differentiation toward most fates, not restricted to fates defined specifically by their rostral/caudal transcription (e.g. caudal mesoderm and caudal neurectoderm), as well as in lineages creating significant caudal-rostral diversity (e.g. neural tube). Yet, rostral/caudal identities of some lineages are inconsistent, possibly reflecting morphological differences. For example, the data indicates strong separation between the large fraction of rabbit extraembryonic mesodermal amnion cells, that develop a strong transient caudal expression signature in E_t_∼7.7, and a minority showing a transient rostral signature, converging after E_t_8 (**Figure S7A**). Similar separation is also seen in the surface ectoderm, though in this case the separation is maintained throughout. The strong expression of the axial gene signature indicating a more polarized development of the amnion in the rabbit vs the mouse, suggests that this module may be used by evolution at different stages and lineages, to compensate for the morphological divergence (size and shape) between the species.

### Screening for evolutionarily diverged differentiation kinetics toward conserved fates

To further characterize the conservation or divergence of gene expression kinetics, we compared entire differentiation trajectories between mouse and rabbit. Specifically, we wished to examine if temporal kinetics of key cell-type-specific genes would be tightly conserved, shifted in phase, or compensated by other genes, and whether the kinetics of genes that are transiently active during state differentiation are also conserved. For example, when studying the differentiation toward somitic mesoderm, we observed high conservation between rabbit and mouse trajectories and expression kinetics (**Figures 6A** and **S7B)**. In-depth analysis of TF kinetics along this trajectory showed that while some key factors are tightly conserved between the species (**Figure 6B**, top), other TFs have altered timing (**Figure 6B**, second row). Regulation of the somitic mesoderm program is also correlated with conserved transient expression kinetics, though such behavior seems to span a more extended period in the mouse (**Figure 6B**, lower rows). Thus, this coordinated network of TFs (such as T, TBX6, and RIPPLY2) not only leads toward the ultimate expression of a conserved somitic mesoderm program (including for example, retinoic acid signaling genes, collagens, etc.), but also to highly conserved timing of the entire somitic mesoderm specification process.

**Figure 6.**
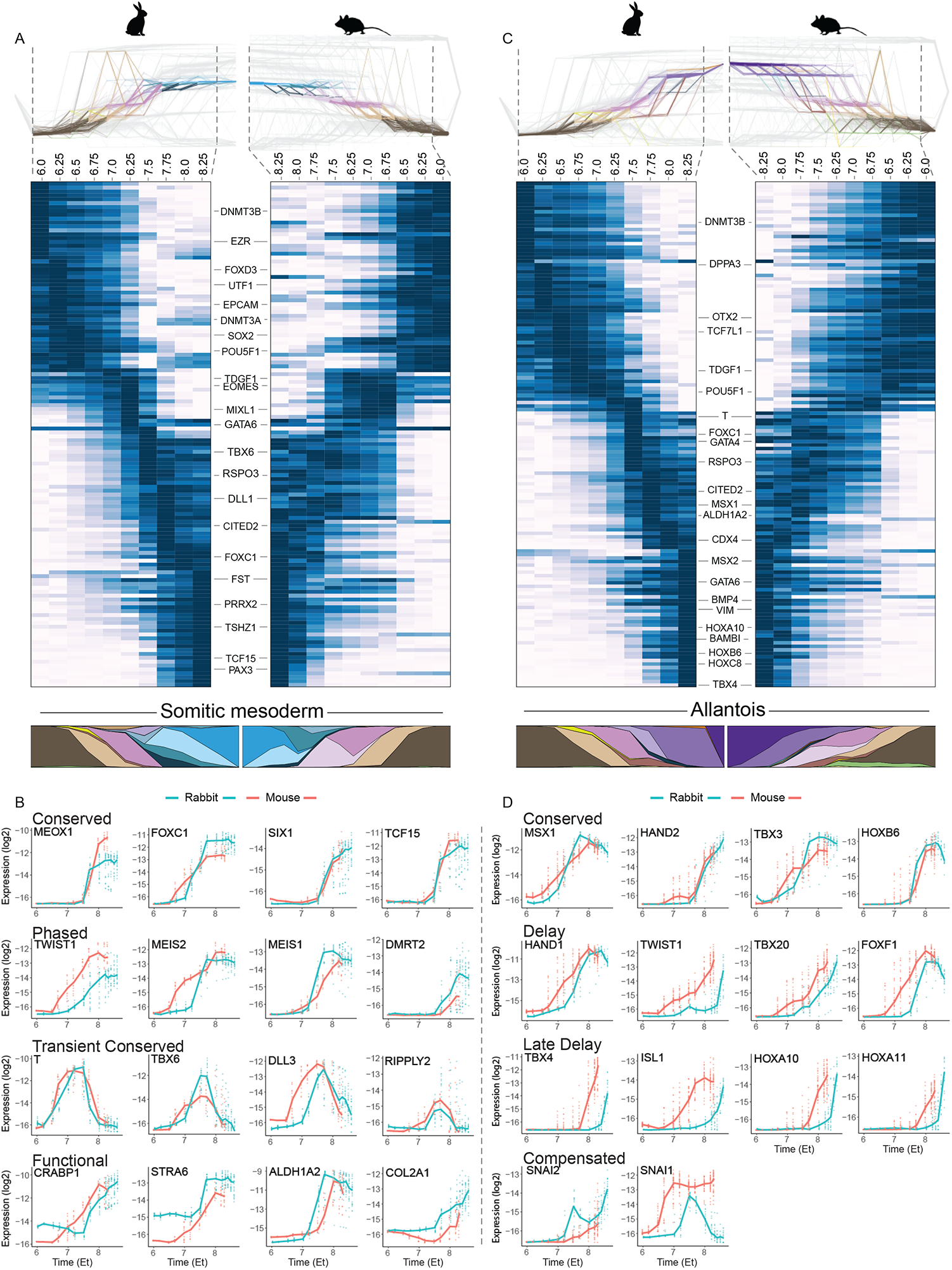
Conservation and divergence in expression kinetics over differentiation tracebacks. (A, B) Tracebacks were performed from (A) somitic-mesoderm and (C) allantois, in flow models of both Rabbit (left) and Mouse (right, mirrored). The latest rabbit time bin, and the first mouse time bin, for which there are no corresponding data from the other species, were trimmed. Network plots depict traced-back flows, colored by cell types (top). Heatmaps show average expression kinetics for genes with conserved gene expression profiles (see Table S3), displaying the highest variance over the trajectories. Below, colored polygons represent the cell type composition of the traceback at each time point for the full flow maps. (B, D) Absolute expression level (log2 of UMI frequency, y-axis) for select marker genes along each of the trajectories shown in the respective network plot. Dots represent individual tracebacks for each terminal MC, lines indicate respective medians.

As a contrary example, the differentiation kinetics of the allantois greatly diverges, and rabbit embryos develop an allantois cell state later and more synchronously than the mouse (**Figures 6C** and **S7C**). Comparative kinetics analysis (**Figure 6D**) outlines the potential mechanistic basis for such divergence. Notably, some TFs that characterize the extraembryonic mesoderm program define a conserved basis for allantois differentiation and emerge at similar time points (**Figure 6D**, top), but other more specific allantois factors are delayed (**Figure 6D** second and third rows). Perhaps most significant is the allantois-specific transcription factor TBX4 which shows a very significant delay in rabbit compared to the mouse. The delayed commitment of the allantois (visible from E9, **Figure S7I**) is possibly related to the aforementioned overall delay in maturation of the rabbit placenta, perhaps relaxing the requirement for its early maturation. In summary, comparing differentiation processes using kinetics of orthologous genes can provide a powerful tool for identifying conserved regulatory links and may suggest how similar regulation is used within diverging contexts. It is however only a basis for even deeper exploration of the evolution of gastrulation logic which must also consider regulatory rewiring events, for example those involving compensation by paralogous genes as shown for SNAI1/2 (**Figure 6D**, bottom), or repurposing factors to perform diverged tasks.

### Early specification of rabbit primordial germ cells without transient mesoderm program activation

From all embryonic gastrulation states, primordial germ cells (PGCs) were associated with much lower conservation scores between the species (**Figures 3A, B**). The flow model indicates that rabbit PGCs appear early at E_t_7.0 (stage3) (**Figure 1C**), and are specified either directly from the epiblast state or from a variant of the epiblast state distinctly marked by TFAP2A and TFAP2C expression (hence denoted “median TFAP”) (**Figure 2A**). In a previous study^7^, numerous TFAP2C/POU5F1 positive cells were observed in the posterior end of gastrulating rabbit embryos. First, broadly in the posterior epiblast layer of stage2 embryos (E6.5-6.75), and later also in the mesoderm layer, becoming more spatially restricted by stage3-4 (E6.5-7.0). Further, in a recent human in vitro study, a TFAP2A/TFAP2C hierarchy was shown to regulate PGC specification from embryonic stem cells (ESCs)^57^. Intriguingly, we find rabbit PGCs rapidly downregulate TFAP2A while TFAP2C levels remain high (**Figures 7A and S8A**), yet a similar cell state does not exist in the mouse (**Figure S8A**). To further examine transcriptional dynamics underlying rabbit PGC specification, we screened for TFs that were specifically enriched in rabbit PGC metacells and compared their time-dependent expression between epiblast, median TFAP, PS, Nascent mesoderm, and PGC. The PGC states are associated with high expression levels of TFAP2C, SOX15, SOX17, CITED1, FOXA3, and NANOS3 (**Figure 7A**). Interestingly, SOX15 has been shown to be a primate and rabbit naïve pluripotency TF, though not in mouse^13,40^. Analysis of PS and nascent mesoderm TFs, clearly showed that PGCs from all embryonic stages do not upregulate most canonical TFs, such as T, EOMES, MIXL1, SNAI1 and HOXA1. This is in contrast to the transient but significant upregulation of the PS genes T and EOMES observed in vitro^7^. In addition, DPPA3 (Stella) that is gradually silenced with time in the epiblast and all mesodermal states (**Figure S8B**) remains expressed throughout in PGCs – further supporting a direct differentiation trajectory. Comparing expression of PGC-specific genes between mouse and rabbit (using epiblast as a common baseline) showed conserved features, including the upregulation of PRDM1, TFAP2C, SOX17, and KIT (**Figure 7B**). Consistent with the previously described downregulation of SOX2 in non-rodent PGCs^7,58^, we find rapid downregulation of this gene in the PGC trajectory starting at the median TFAP progenitor state (**Figure S8C**). The expression of Fragilis (IFITM3), ALPL, and additional mouse-specific genes were completely missing in rabbit PGC (some possibly due to technical lack of orthology). At the same time, a number of the strongest rabbit markers were either not expressed in mouse (CD34, RELN, FOXA3), or did not have a mouse orthologue, such as the rabbit orthologue for human germ-cell-specific gene PASD1^59^. The early specification of rabbit PGCs is also supported by cell cycle analysis (**Figure 7C)**. While it is impossible to accurately deduce the actual reduction in PGC proliferation, the data suggest it to be mild, and stable up to E_t_8.6. Together with flow analysis (**Figure S8D**), these results are most consistent with an early specified self-sustaining PGC population with declining frequency but slowly increasing absolute numbers, within a more rapidly growing embryo.

**Figure 7.**
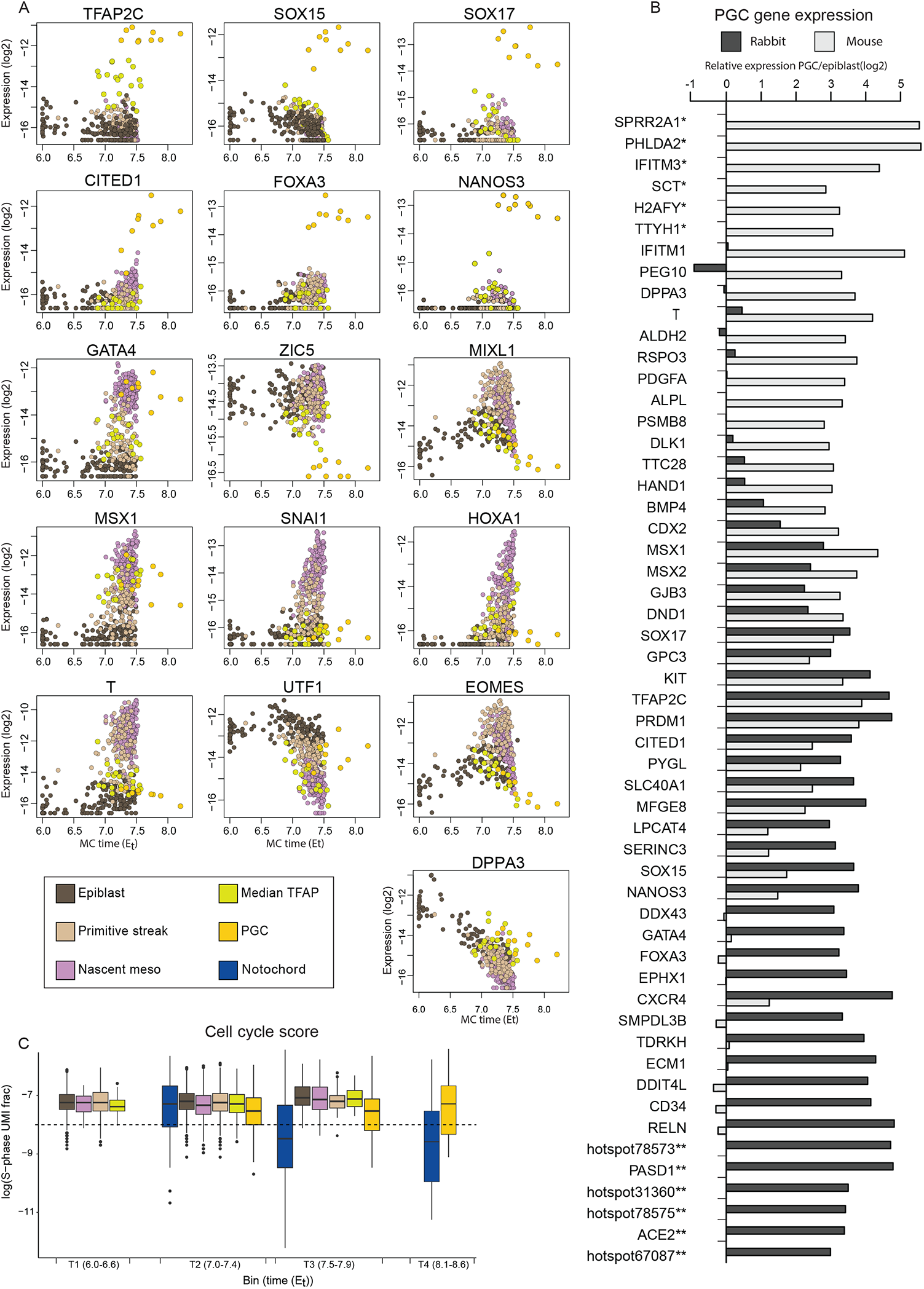
PGC differentiation. (A) Absolute expression level (log2 of UMI frequency, y axis) for select marker genes. (B) Relative expression in PGC vs epiblast (log2 of UMI frequency in PGC - log2 of UMI frequency in epiblast) in rabbit (Dark grey) and mouse (light grey). Mouse genes with no known rabbit ortholog are annotated with “*” (top) and rabbit genes lacking mouse orthology with “**” (bottom). (C) Cell cycle score. A cycling score combining the expression of S- and M-phase characteristic genes (see Methods) was assigned to each single cell (y-axis). Due to the low number of PGCs in the manifold, cells were binned to four groups (6-6.6, 7-7.4, 7.5-7.9, 8.1-8.6). Partial cycling slowdown in the PGC population is observed, when compared to the notochord (substantially halting) or other early populations (fully cycling).

## DISCUSSION

Gastrulation is the hallmark developmental process that translates “raw” intracellular pluripotent regulatory logic into the stereotypical blueprint of the vertebrate body plan. It requires single cells to acquire increasingly more specialized internal states in a precise and stable fashion. At the same time, and perhaps more importantly, gastrulation is defined by the capacity of cell ensembles to acquire such identity in a coordinated fashion. All the while, the embryo proliferates massively, and the initially unlimited potential of the epiblast is restricted and shaped by a combination of changes in cells’ internal state, cellular interactions, and physical forces - toward defined germ layers, cell types, and organ precursors. In this work, we used extensive sampling of the ensemble of single cells from single gastrulating rabbit embryos to combine quantitative characterization of transcriptional cell states with a model tracking the coordinated dynamics of the cell ensembles over time. Using comparative analysis of the new rabbit model and our recently published mouse gastrulation model^14^, we could begin to approach fundamental questions on the universal and conserved properties of mammalian gastrulation and characterize diverged mechanisms ultimately leading toward the diversity across mammals in general, and between rabbit and mouse in particular.

An initial “static” comparative strategy of mouse and rabbit gastrulation manifolds shows remarkable conservation between the transcriptional states of both species. Such conservation is both specific and quantitative. Its specificity is indicated by the identification of 79 distinct RBOMs - transcriptional states that are reciprocally matched between the species. The discovery of a large number of distinct RBOMs reflects a truly rich conserved transcriptional space going far beyond a broad conservation of the germ layers states or specialized cell types like the primitive erythrocytes. Furthermore, conservation is highly quantitative, as shown by comparison of gene expression frequencies of thousands of genes in matched states, indicating almost all genes are conserved quantitatively up to a two-fold ratio. This precision of conservation is surprising since the gastrulation manifold does not consist of a large number of highly distinct states, but is instead a continuum of multifurcating programs^14,23^. Such plasticity could in theory support significant regulatory redundancy and subsequent evolutionary flexibility. But instead, at the core of the gastrulation programs, which we define by reciprocal best orthologous genes, we can identify 76 (and possibly more) transcription factors that retain a distinct and rigid regulatory identity between mouse and rabbit. Such conservation suggests that these factors coordinate the gastrulation process in a highly quantitative and balanced fashion, which limits the evolutionary plasticity of this process. Notably, 7 of these TFs are of the HOX family, supporting their suggested central role in establishing the phylotypic state^1^. Our model therefore calls for a quantitative and multi-factorial approach toward understanding the regulation of gastrulation state transitions in ways that were so far difficult to practice.

As noted above, understanding gastrulation at the transcriptional level entails more than mapping static cell states and types at single cell resolution, as the process inherently involves ensembles of interactive and proliferating cells. We therefore developed a framework for comparative analysis of the gastrulation process as it occurs over a carefully quantified time scale. The data coming out of this analysis unequivocally showed that gastrulation is conserved as a process, and highlighted a specific point in time, coinciding with the onset of massive differentiation (stages 4c-5; E_t_7.5-7.7), where both mouse and rabbit gastrulation are aligned with maximum specificity (**Figure 5A**). This striking conservation, which provides fresh support and a possible mechanistic basis to the hourglass model for vertebrate evolution should be considered in light of the highly diverged extraembryonic tissues, overall difference in size, as well as dramatic morphological differences between the rabbit bilaminar disc and mouse cup-shaped structures^4^. Still, the structural divergence of the rabbit and mouse extraembryonic endoderm remains compatible with conserved signaling. Namely, the canonical organizer function of the anterior hypoblast and the corresponding mouse AVE. One key difference however, is the apparent compensation by the rabbit hypoblast for the lack of functional signaling from the trophoblast. In general, rabbit trophoblast is markedly delayed in its development compared to the mouse extraembryonic ectoderm, in line with similar retardation of the allantois. The divergence of extraembryonic structures are contradictory to the strong conservation of the gastrulation process. This could suggest that the crucial conserved steps underlying embryonic diversification are more intrinsically driven, for example by the rich core of conserved TFs discussed above. Such a process is unlikely to rely on complex maternal or extraembryonic interactions that would strongly depend on conserved structure.

Our detailed rabbit gastrulation flow model and the new comparative methodology hereby provide a scaffold for further comprehensive and robust interpretation of other mammalian models, for which similar resolution and accuracy is yet to be attained. Human (and primate) gastrulation is already known to share many states and characteristics with the mouse. However, the temporal process of differentiation and proliferation for human cell ensembles during gastrulation would have to be modeled more carefully, especially considering the sparsity of such data. As flow models become integrated with new data and models for primates and other species, it can be expected that we will come closer toward truer understanding of human developmental process. This, will be useful for designing the control of embryonic stem cell differentiation, tissue models, and analysis of ex-utero and synthetic embryos. Lastly, such models will place us in an unprecedented position to synthesize the molecular phenotypes of gastrulation states between and within phyla^60^ with an underlying genome and epigenome evolutionary theory.

## FIGURE LEGENDS

Figure S1. **Rabbit embryo compendium**

Pictures of embryos sampled in the study. Embryos shown in sequence according to intrinsic k-NN transcriptional rank. Color bars indicates separation into 12 age groups. Scale bar = 500um.

Figure S2. **Embryo collection and scRNA-seq data preparation**

(A) Distribution of UMIs per single cell, exclusion cells criteria denoted by red and green vertical lines.

(B) Distribution of fraction of excluded gene UMIs per single cell (see Metacells vignette). Exclusion cells criteria denoted by red vertical line.

(C) Number of embryonic cells per embryo (after QC, ranked from “youngest” to “oldest”).

(D) Number of embryos participating in the flow analysis, collected over gestation time.

(E) Embryo stages by morphological assessment vs gestation time.

(F) Calculated time (Et) vs gestation time for each time bin.

(G) Genome annotation enhancement pipeline.

(H) Temporal ordering and binning of embryos. Left, k-NN embryo/embryo similarity (horizontal lines mark separation of time bins, numbered in bold). Right, cell-type frequency per embryo (see Figure 1).

Figure S3. **Flows support**

(A) Embryonic cells per time bin, plotted over the 2-D UMAP projection, color coded by cell type.

(B) MC/MC distance metric used as constraint for the flow problem formulation, indicating transcriptional logistic distance.

(C) Cell cycle S-phase score vs M-phase score based on UMI count fractions. Cutoff line (bottom left diagonal) represents cells with low cycling score.

(D) Distribution of MCs cycling score, by type. MC cycling score indicates fraction of cells below cutoff (as in C).

Figure S4. **MC alignment support**

(A) Correlation across all MCs of the rabbit and mouse manifolds, ordered by cell type and internally by time.

(B) Correlation across the 79 RBOMs of the rabbit and mouse manifolds (ordered as in (A)).

(C) Example of transcriptionally diverged sequence orthologs of APLN.

(D) Example of sequence orthologs showing species-specific constant bias. Shown before and after linear correction (factor of 0.41) according to the manifold alignment.

(E) Example of new putative ortholog association for un-annotated rabbit MESP1 gene, matched with the transcriptional hostpot33433.

(F) Example of annotation assignment for the rabbit TWIST1 gene based on alignment to the mouse genome (mm9). Table lists explicit OryCun3.0 genome coordinates for genes mentioned in (C-F).

Figure S5. **Extraembryonic ectoderm (ExEc) interspecies comparison**

(A) Side by side gene expression heatmap in mouse and rabbit ExEc.

(B) MC-MC correlation matrix of rabbit and mouse ExEc.

(C) Relative expression of notable ExEc and ExEnd genes (rabbit only).

(D) Temporal distributions of rabbit ExEc.

(E) Temporal expression (log2 of UMI frequency) of key genes during ExEc development. Each dot represents a single MC.

(F, G) Temporal expression of rabbit BMP4 and mouse IGF1, respectively. Each dot represents a single MC.

Figure S6. **Conserved and diverged gene expression signatures in the epiblast and PS**

(A) Comparison of PS - Epiblast differential gene expression (log2) between rabbit (x-axis) and mouse (y-axis). Canonical TFs such as MIXL1 and T are upregulated in the epiblast to PS transition in both species (top right corner).

(B) Heatmap of genes with extreme profiles in their PS-Epiblast differential expression, grouped by the quadrants of (A), each column represents a single MC.

Figure S7. **Gene expression trends over trajectories**

(A-H) Rostral-Caudal score of individual MC trajectories of terminal types, as indicated.

(I) Representative E9.0 embryo (16 somite pairs, not sequenced).

Figure S8. **PGC specification**

(A) Expression over time of TFAP2A for mouse and rabbit.

(B) Expression of DPPA3 (Stella) over time. While gradually silenced with time in the epiblast and mesodermal states, DPPA3 remains expressed in PGCs.

(C) Expression over time of SOX2 for mouse and rabbit.

(D) Internal PGC flow over time. After an initial differentiation of non-PGC cell types to PGC, beyond Et7, the number of PGC cells can be explained solely by internal proliferation.

## METHODS

### Embryo Collection and Documentation

Embryos were collected from pregnant New Zealand White female rabbits (mated by Envigo, and shipped on the morning of each experiment), between E6.0-8.5. All animal experiments were approved by the Institutional Animal Care and Use Committee (07100820-2). Animals were sacrificed by intravenous injection of lethal dose of pentobarbital (100mg/kg), and embryos removed as described by^61^. In brief, uteri were placed in ice-cold PBS, and embryos removed individually from dissected uterus segments containing a single site using fine iridectomy scissors on darkened (Activated charcoal C9157, Sigma-Aldrich) Sylgard184 (Dow) coated 10cm plates. Embryos were resected from most of the extraembryonic tissues, washed in fresh PBS, and transferred to chilled DMEM (Phenol-red free, Gibco) supplemented with 10% FBS (Biological Industries) for imaging prior to dissociation. Phase contrast images were taken with an Eclipse Ti2 inverted microscope (Nikon) and Zyla sCMOS camera (Andor). Embryo staging was performed according to^36,62^.

### Flow Cytometry

Prior to dissociation, to focus on tissues that contribute to the embryo proper, most of the extraembryonic ectoderm and yolk sac were removed using fine forceps and fine iridectomy scissors. For isolation of single cells for scRNA-seq, embryos were dissociated with 0.25% Trypsin-A, 0.02% EDTA (Biological Industries) solution for 5’ at 37°C, and resuspended in DMEM w/o phenol red (Gibco) supplemented with 10% FBS (Biological Industries). Samples were sorted to 384well lysis plates using a FACSAria-III or Symphony S6 flow cytometer (BD Biosciences), excluding doublets and cell debris by FSC and SSC metrics.

### Single-cell RNA-sequencing and initial filtering

Single cell cDNA libraries were prepared using the MARS-seq method, as described^35,63^, with the following modifications: The final concentration of the RT1 primers was 2nM, and pooling was done via centrifugation to VBLOCK200 reservoir (Clickbio). Klenow reaction was not followed by heat inactivation. The volume of the first RT and Exonuclease I reactions mix were scaled down to 1 and 0.5ul, respectively, and dispensed by MANTIS liquid handler (FORMULATRIX). MARS-seq reads were processed following the MARS-seq2.0 pipeline^63^. Briefly, demultiplexing plate barcodes (4 base pairs) from Read 1, the reads were mapped to the OryCun3.0 genome using STAR^64^. Demultiplexing of the reads and construction of the single-cell UMI matrix was based on well barcodes (7 base pairs) and UMI barcodes (8 base pairs) from Read 2. Embryo identities were recorded per well during sorting and recovered as part of the MARS-seq post-processing stage. Overall we processed 165,504 wells, out of which we retained 117,660 well-covered cells for further analysis. In particular, cells with less than 2,600 or over 50,000 UMIs were filtered out (median 7,142 UMIs).

### Reference genome and annotation

We based our annotated 3’ gene ends on the rabbit reference genome OryCun2.0 [GCA_000003625.1]. This assembly is of shallow depth at 7.5X and annotated with limited precision. We’ve extended the basic annotation with additional RefSeq gene orthologies to define a total of 48,189 genes. We then used the higher depth (but not annotated) OryCun3.0 assembly [GCA_013371645.1], and projected the initial set of OryCun2 annotations to it using LiftOver mapping. This resulted in 42,926 genes being uniquely lifted. We expected this set of predicted 3’ends to be missing genes in poorly aligned genomic regions and therefore performed additional screening for genes using clusters of mapped of scRNA-seq reads that could not be matched with an existing annotated region. This was done by identifying *transcriptional hotspots* as genomic bins of 500 bps that were covered by over 150 scRNA-seq UMIs across our dataset, unifying overlapping bins. Hotspots that ended up overlapping with an annotated 3’ locus were merged into the annotated gene model. For hotspots that were not associated with genes in this way, we attempted additional annotation using blastn with the refseq_rna db. A summary of the annotation pipeline is shown in **Fig S2G**. The extended annotation is available from the papers’ github companion.

### Rabbit embryo k-NN ordering

Following an initial coarse-grained embryo ordering, we applied the k-NN order refinement heuristic as we previously described for the mouse^14^. Briefly, a cell-cell transcriptional similarity k-NN graph is constructed over all embryos’ cells, excluding those annotated as extraembryonic. Embryo similarity is defined by the number of k-NN edges crossing between the embryos’ cells, normalized for the number of cells per embryo. A greedy pairwise order switching heuristic aims at eliminating k-NN edges crossing large time gaps, improving upon the initial ordering until convergence (**Figure S2H**).

### Defining absolute developmental time

Each embryo was associated with an approximate time based on its sampling time post mating. For the latest somite stages, an idealized time was given based on a 2hr cycle for each somite (see **Table S1** for data on all embryos). We then formed 12 bins aiming to equalize the total number of cells per bin and its time homogeneity. We then manually assigned a representative time per bin, defining the parameter 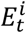 for bin i (**Figure 1C, S2H**).

### Metacell analysis and cell type annotation

Metacells^37^ analysis was performed as described in the package vignette. Mitochondrial associated hotspot transcripts and additional genes linked with non-coding activities that are batch-affected were excluded, removing 391 loci in total. A total of 1,565 stress, cell cycle-related, and additional batch-affected transcripts in manually annotated gene modules were defined as forbidden to be features prior to applying metacells. Metacells then derived a metacell model over 3,584 MCs representing 117,660 retained single cell profiles.

### Cell cycle modules and estimation of metacell proliferation rate

We defined the M-phase gene module using single cell correlation to MKI67, TOP2A, and UBE2C, and expanded to include by association ARHGAP11A, ARHGAP27P1-BPTFP1-KPNA2P3, ARL6IP1, ASPM, AURKA, AURKB, BUB3, CCNB1, CCNB2, CCNF, CDCA2, CDCA3, CENPE, CENPF, CKAP2, CKAP2L, DEPDC1B, G2E3, GTSE1, HYLS1, INCENP, KIF11, KIF14, KIF18A, KIF20A, KIF23, KIFC1, MIS18BP1, MKI67, NCAPG, NUSAP1, PLK1, PRC1, PRR11, PSRC1, RACGAP1, SGO2, SPAG5, TOP2A, TPX2, TTK, UBE2C and hotspot44564.

The group of S-phase genes was defined by correlation to PCNA, RRM2 and expanded to include DTL, FEN1, MCM2, MCM4, MCM6, MCM7, PCNA, RFC3, RRM2, hotspot60823.

For each cell, we computed an M- and S-phase score as the fraction of UMIs for these gene modules (**Figure S3C**). Based on these estimations we assigned each metacell m with a proliferation rate in units of doublings per 24h:

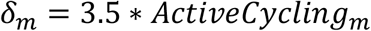

Where 3.5 is the maximum doublings per day, and *ActivitCycling*_*m*_ is defined as the fraction of cells showing high combined S- and M-phase expression in metacell m:

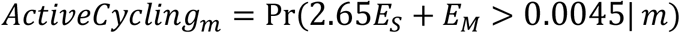

### Network flow inference and modeling

We inferred rabbit network flows as previously described for the mouse^14^, with improved modelling of proliferation rate variation among metacells. The standard network flow constraint is assuming all nodes (e.g. metacells) are conserving their relative frequencies, which can be interpreted as either constant mass per time, or as assuming uniform proliferation rates:

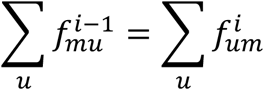

Where m, u are metacells, i the time and f the flow per time and metacell pair.

To express variable proliferation rate per metacell, we define generalized flow constraints as:

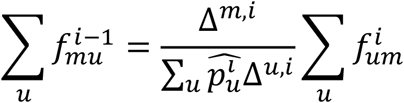

Where the growth per metacell and time Δ is defined as:

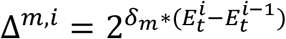

And 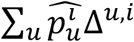 represent the mean growth of the ensemble in time bin i, with 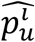 is defining the estimated frequency of metacell u at time bin i.

We inferred *δ* values for all metacells, except for those annotated as notochord and gut for which *δ*_*m*_ = 0 and *δ*_*m*_ = 1 were assumed based on empirical analysis of overall cell type frequency over time, respectively. In addition, for hematopoietic lineages we defined negative proliferation rates to account for their under sampling. This, due to the radial expansion of these cells in the extraembryonic yolk sac mesoderm, that results in their progressive elimination during dissection.

### Manifold alignment

To facilitate comparison of transcriptional states between rabbit and mouse, we first identified 9,748 gene pairs with common names in the two species, representing a set of genes G with unambiguous sequence orthologies for downstream analysis. We then represented the rabbit and mouse manifolds quantitatively using distributions of gene expression for this set of comparable genes over metacells, log transformed and regularized (*log*2(10^−5^ + *p*)) to defined matrices 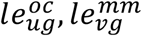 for rabbit (oc) and mouse (mm) respectively. We then compared metacells U and V in the two species using Pearson correlation of the log transformed vectors to derive:

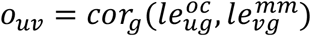

We then identified reciprocally best orthologous metacells (RBOMs, **Figure S4B**) pairs as:

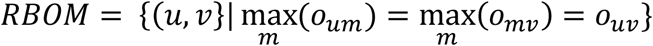

That is, for each metacell in one manifold, we picked the maximal correlating metacell in the other manifold only if the correlation was also maximal for that second manifold metacell across all the metacells of the first. Using the RBOMs as a comparable manifold reference, we computed gene-gene cross-species correlations:

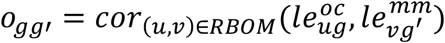

To correct for systematic estimation biases between the species we computed a linear model (setting constant to zero) fitting each rabbit gene expression (*in linear scale*: 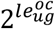) to its orthologous mouse genes over the RBOMs (**Fig S4D**). Whenever the best fit coefficient was in the range 0.2-5 we applied it to correct rabbit expression, and used corrected values for the comparative work described in Fig 5-7.

### Projected annotation

We used the correlation between corrected expression profiles of the rabbit manifold and the mouse expression profiles to redefine the similarities between all rabbit and mouse metacells (RBOM being the subset for which matching is completely unambiguous) (**Fig S4A**). To allow comparisons of whole embryos between the two species, we represented mouse embryos as distribution over the mouse cell type annotation terms. We then derived a similar representation (over the same mouse types) for rabbit embryos in an unbiased fashion. To this end, we identified the top 5 correlated mouse metacells for each rabbit metacell, and selected the majority annotation term within them as the assigned projected rabbit annotation term. This provided us with a common representation for embryos in the two species, allowing computation of the cross-species embryo correlation as shown in **Fig 5A**. We linearly interpolated the cell type frequencies per time bins and censored it at Et6-8.3, to create a high-resolution common time grid for both species. To derive the trend line shown in **Fig 5A**, we used dynamic time warping (DTW) applied on the two time series.

### Inference of kinetics over genes and gene modules

To infer the differentiation trajectories leading to cell fates of interest (**Figure 5G, Figure 6, Figure S7**), we used the flow model as described in Mittnenzweig et al 2021^14^. When analyzing the kinetics of gene modules (e.g., Rostral/Caudal analysis in Figure 5), we used the log2 transformed mean of the expression per metacell over the module’s genes (eg. Rostral: CER1, OTX1, SHISA2, HESX1, and Caudal: GBX2, TBX6, CDX1,2,4, GRSF1).

## ACKNOWLEDGMENTS

We thank the Tanay and Stelzer group members for discussion and advice. Special thanks to Hernan Rubinstein for help in graphic design and embryo illustrations. Y.S. is the incumbent of the Louis and Ida Rich Career Development Chair and a member of the European Molecular Biology Organization (EMBO) Young Investigator Program. Research in the Stelzer lab is supported by European Research Council (ERC_StG 852865), Moross Integrated Cancer Center, Helen and Martin Kimmel Stem Cell Institute, the Schwartz/Reisman Collaborative Science Program, and Abisch Frenkel Foundation. This research was also generously supported by Barry and Janet Lang, Hadar Impact Fund, Lord Sieff of Brimpton Memorial Fund, Janet and Steven Anixter, JoAnne Silva, Maurice and Vivienne Wohl Biology Endowment, and Lester and Edward Anixter Family. M.M. was a postdoctoral fellow of the Minerva Stiftung and is supported by the Walter Benjamin program of the German Research Foundation (DFG). A.T. is supported by the European Research Council (ERC CoG scAssembly), the EU BRAINTIME project, the Israel Science Foundation, and the Chen-Zuckerberg Foundation. This research was further supported Israeli Council for Higher Education (CHE) Data Science program, and by a grant from Madame Olga Klein-Astracha.

## AUTHOR CONTRIBUTIONS

Y.M., O.R., A.T., and Y.S. conceived and designed the experiment, and performed data analysis and its interpretation. S.C., R.B.Y, and R.H. assisted in the collection of single-cells from individual embryos. N.R. prepared scRNA-seq libraries. A.L., M.M., and O.B.K. assisted with scRNA-seq data analyses. Y.M., O.R., A.T., and Y.S. wrote the manuscript with input from all the authors.

## REFERENCES

1. Duboule, D. (1994). Temporal colinearity and the phylotypic progression: a basis for the stability of a vertebrate Bauplan and the evolution of morphologies through heterochrony. Dev Suppl, 135–142.

2. Raff, R.A. (1996). The Shape of Life: Genes, Development, and the Evolution of Animal Form (University of Chicago press).

3. Domazet-Loso, T., and Tautz, D. (2010). A phylogenetically based transcriptome age index mirrors ontogenetic divergence patterns. Nature 468, 815–818. 10.1038/nature09632.

4. Eakin, G.S., and Behringer, R.R. (2004). Diversity of germ layer and axis formation among mammals. Semin Cell Dev Biol 15, 619–629. 10.1016/j.semcdb.2004.04.008.

5. King, B.F., and Enders, A.C. (1993). Comparative Development of the Mammalian Yolk Sac. In The Human Yolk Sac and Yolk Sac Tumors, F.F. Nogales, ed. (Springer Berlin, Heidelberg), pp. 1–32. 10.1007/978-3-642-77852-0.

6. Irie, N., Tang, W.W., and Azim Surani, M. (2014). Germ cell specification and pluripotency in mammals: a perspective from early embryogenesis. Reprod Med Biol 13, 203–215. 10.1007/s12522-014-0184-2.

7. Kobayashi, T., Castillo-Venzor, A., Penfold, C.A., Morgan, M., Mizuno, N., Tang, W.W.C., Osada, Y., Hirao, M., Yoshida, F., Sato, H., et al. (2021). Tracing the emergence of primordial germ cells from bilaminar disc rabbit embryos and pluripotent stem cells. Cell Rep 37, 109812. 10.1016/j.celrep.2021.109812.

8. Denker, H.W. (2000). Structural dynamics and function of early embryonic coats. Cells Tissues Organs 166, 180–207. 10.1159/000016732.

9. Fischer, B., Chavatte-Palmer, P., Viebahn, C., Navarrete Santos, A., and Duranthon, V. (2012). Rabbit as a reproductive model for human health. Reproduction 144, 1–10. 10.1530/REP-12-0091.

10. Viebahn, C., Mayer, B., and Hrabe de Angelis, M. (1995). Signs of the principle body axes prior to primitive streak formation in the rabbit embryo. Anat Embryol (Berl) 192, 159–169. 10.1007/BF00186004.

11. Nishimura, M. (2001). Timing of implantation in New Zealand White rabbits. Congenital Anomalies, 198–203.

12. Argelaguet, R., Clark, S.J., Mohammed, H., Stapel, L.C., Krueger, C., Kapourani, C.A., Imaz-Rosshandler, I., Lohoff, T., Xiang, Y., Hanna, C.W., et al. (2019). Multi-omics profiling of mouse gastrulation at single-cell resolution. Nature 576, 487–491. 10.1038/s41586-019-1825-8.

13. Bouchereau, W., Jouneau, L., Archilla, C., Aksoy, I., Moulin, A., Daniel, N., Peynot, N., Calderari, S., Joly, T., Godet, M., et al. (2022). Major transcriptomic, epigenetic and metabolic changes underlie the pluripotency continuum in rabbit preimplantation embryos. Development 149. 10.1242/dev.200538.

14. Mittnenzweig, M., Mayshar, Y., Cheng, S., Ben-Yair, R., Hadas, R., Rais, Y., Chomsky, E., Reines, N., Uzonyi, A., Lumerman, L., et al. (2021). A single-embryo, single-cell time-resolved model for mouse gastrulation. Cell 184, 2825–2842 e2822. 10.1016/j.cell.2021.04.004.

15. Niu, Y., Sun, N., Li, C., Lei, Y., Huang, Z., Wu, J., Si, C., Dai, X., Liu, C., Wei, J., et al. (2019). Dissecting primate early post-implantation development using long-term in vitro embryo culture. Science 366. 10.1126/science.aaw5754.

16. Nowotschin, S., Setty, M., Kuo, Y.Y., Liu, V., Garg, V., Sharma, R., Simon, C.S., Saiz, N., Gardner, R., Boutet, S.C., et al. (2019). The emergent landscape of the mouse gut endoderm at single-cell resolution. Nature 569, 361–367. 10.1038/s41586-019-1127-1.

17. Pijuan-Sala, B., Griffiths, J.A., Guibentif, C., Hiscock, T.W., Jawaid, W., Calero-Nieto, F.J., Mulas, C., Ibarra-Soria, X., Tyser, R.C.V., Ho, D.L.L., et al. (2019). A single-cell molecular map of mouse gastrulation and early organogenesis. Nature 566, 490–495. 10.1038/s41586-019-0933-9.

18. Rayon, T., Maizels, R.J., Barrington, C., and Briscoe, J. (2021). Single-cell transcriptome profiling of the human developing spinal cord reveals a conserved genetic programme with human-specific features. Development 148. 10.1242/dev.199711.

19. Ton, M.-L.N., Keitley, D., Theeuwes, B., Guibentif, C., Ahnfelt-Rønne, J., Andreassen, T.K., Calero-Nieto, F.J., Imaz-Rosshandler, I., Pijuan-Sala, B., Nichols, J., et al. (2022). Rabbit Development as a Model for Single Cell Comparative Genomics. bioRxiv.

20. Tyser, R.C.V., Mahammadov, E., Nakanoh, S., Vallier, L., Scialdone, A., and Srinivas, S. (2021). Single-cell transcriptomic characterization of a gastrulating human embryo. Nature 600, 285–289. 10.1038/s41586-021-04158-y.

21. Wagner, D.E., Weinreb, C., Collins, Z.M., Briggs, J.A., Megason, S.G., and Klein, A.M. (2018). Single-cell mapping of gene expression landscapes and lineage in the zebrafish embryo. Science 360, 981–987. 10.1126/science.aar4362.

22. Qiu, C., Cao, J., Martin, B.K., Li, T., Welsh, I.C., Srivatsan, S., Huang, X., Calderon, D., Noble, W.S., Disteche, C.M., et al. (2022). Systematic reconstruction of cellular trajectories across mouse embryogenesis. Nat Genet 54, 328–341. 10.1038/s41588-022-01018-x.

23. Moris, N., Pina, C., and Arias, A.M. (2016). Transition states and cell fate decisions in epigenetic landscapes. Nat Rev Genet 17, 693–703. 10.1038/nrg.2016.98.

24. Tritschler, S., Buttner, M., Fischer, D.S., Lange, M., Bergen, V., Lickert, H., and Theis, F.J. (2019). Concepts and limitations for learning developmental trajectories from single cell genomics. Development 146. 10.1242/dev.170506.

25. Cheng, S., Mittnenzweig, M., Mayshar, Y., Lifshitz, A., Dunjic, M., Rais, Y., Ben-Yair, R., Gehrs, S., Chomsky, E., Mukamel, Z., et al. (2022). The intrinsic and extrinsic effects of TET proteins during gastrulation. Cell 185, 3169–3185 e3120. 10.1016/j.cell.2022.06.049.

26. Sankaran, V.G., Weissman, J.S., and Zon, L.I. (2022). Cellular barcoding to decipher clonal dynamics in disease. Science 378, eabm5874. 10.1126/science.abm5874.

27. Chen, A., Liao, S., Cheng, M., Ma, K., Wu, L., Lai, Y., Qiu, X., Yang, J., Xu, J., Hao, S., et al. (2022). Spatiotemporal transcriptomic atlas of mouse organogenesis using DNA nanoball-patterned arrays. Cell 185, 1777–1792 e1721. 10.1016/j.cell.2022.04.003.

28. Liu, C., Li, R., Li, Y., Lin, X., Zhao, K., Liu, Q., Wang, S., Yang, X., Shi, X., Ma, Y., et al. (2022). Spatiotemporal mapping of gene expression landscapes and developmental trajectories during zebrafish embryogenesis. Dev Cell 57, 1284–1298 e1285. 10.1016/j.devcel.2022.04.009.

29. Peng, G., Suo, S., Cui, G., Yu, F., Wang, R., Chen, J., Chen, S., Liu, Z., Chen, G., Qian, Y., et al. (2019). Molecular architecture of lineage allocation and tissue organization in early mouse embryo. Nature 572, 528–532. 10.1038/s41586-019-1469-8.

30. Nakamura, T., Okamoto, I., Sasaki, K., Yabuta, Y., Iwatani, C., Tsuchiya, H., Seita, Y., Nakamura, S., Yamamoto, T., and Saitou, M. (2016). A developmental coordinate of pluripotency among mice, monkeys and humans. Nature 537, 57–62. 10.1038/nature19096.

31. Nakamura, T., Yabuta, Y., Okamoto, I., Sasaki, K., Iwatani, C., Tsuchiya, H., and Saitou, M. (2017). Single-cell transcriptome of early embryos and cultured embryonic stem cells of cynomolgus monkeys. Sci Data 4, 170067. 10.1038/sdata.2017.67.

32. Xu, Y., Zhang, T., Zhou, Q., Hu, M., Qi, Y., Xue, Y., Wang, L., Nie, Y., Bao, Z., and Shi, W. (2022). A single-cell transcriptome atlas of human early embryogenesis. bioRxiv.

33. Ihara, T. (1970). Comparative study of developmental progress in the mouse, rat and rabbit in their stages of organogenesis. Congenital Anomalies 10 (2), 67–81.

34. Williams, B.S., and Biggers, J.D. (1990). Polar trophoblast (Rauber’s layer) of the rabbit blastocyst. Anat Rec 227, 211–222. 10.1002/ar.1092270210.

35. Jaitin, D.A., Kenigsberg, E., Keren-Shaul, H., Elefant, N., Paul, F., Zaretsky, I., Mildner, A., Cohen, N., Jung, S., Tanay, A., and Amit, I. (2014). Massively parallel single-cell RNA-seq for marker-free decomposition of tissues into cell types. Science 343, 776–779. 10.1126/science.1247651.

36. Blum, M., Andre, P., Muders, K., Schweickert, A., Fischer, A., Bitzer, E., Bogusch, S., Beyer, T., van Straaten, H.W., and Viebahn, C. (2007). Ciliation and gene expression distinguish between node and posterior notochord in the mammalian embryo. Differentiation 75, 133–146. 10.1111/j.1432-0436.2006.00124.x.

37. Ben-Kiki, O., Bercovich, A., Lifshitz, A., and Tanay, A. (2022). Metacell-2: a divide-and-conquer metacell algorithm for scalable scRNA-seq analysis. Genome Biol 23, 100. 10.1186/s13059-022-02667-1.

38. Sebe-Pedros, A., Saudemont, B., Chomsky, E., Plessier, F., Mailhe, M.P., Renno, J., Loe-Mie, Y., Lifshitz, A., Mukamel, Z., Schmutz, S., et al. (2018). Cnidarian Cell Type Diversity and Regulation Revealed by Whole-Organism Single-Cell RNA-Seq. Cell 173, 1520–1534 e1520. 10.1016/j.cell.2018.05.019.

39. Tanay, A., and Sebe-Pedros, A. (2021). Evolutionary cell type mapping with single-cell genomics. Trends Genet 37, 919–932. 10.1016/j.tig.2021.04.008.

40. Boroviak, T., Stirparo, G.G., Dietmann, S., Hernando-Herraez, I., Mohammed, H., Reik, W., Smith, A., Sasaki, E., Nichols, J., and Bertone, P. (2018). Single cell transcriptome analysis of human, marmoset and mouse embryos reveals common and divergent features of preimplantation development. Development 145. 10.1242/dev.167833.

41. Hodge, R.D., Bakken, T.E., Miller, J.A., Smith, K.A., Barkan, E.R., Graybuck, L.T., Close, J.L., Long, B., Johansen, N., Penn, O., et al. (2019). Conserved cell types with divergent features in human versus mouse cortex. Nature 573, 61–68. 10.1038/s41586-019-1506-7.

42. Arnold, S.J., and Robertson, E.J. (2009). Making a commitment: cell lineage allocation and axis patterning in the early mouse embryo. Nat Rev Mol Cell Biol 10, 91–103. 10.1038/nrm2618.

43. Cindrova-Davies, T., Jauniaux, E., Elliot, M.G., Gong, S., Burton, G.J., and Charnock-Jones, D.S. (2017). RNA-seq reveals conservation of function among the yolk sacs of human, mouse, and chicken. Proc Natl Acad Sci U S A 114, E4753–E4761. 10.1073/pnas.1702560114.

44. Hoshino, H., Shioi, G., and Aizawa, S. (2015). AVE protein expression and visceral endoderm cell behavior during anterior-posterior axis formation in mouse embryos: Asymmetry in OTX2 and DKK1 expression. Dev Biol 402, 175–191. 10.1016/j.ydbio.2015.03.023.

45. Robb, L., and Tam, P.P. (2004). Gastrula organiser and embryonic patterning in the mouse. Semin Cell Dev Biol 15, 543–554. 10.1016/j.semcdb.2004.04.005.

46. Yoshida, M., Kajikawa, E., Kurokawa, D., Tokunaga, T., Onishi, A., Yonemura, S., Kobayashi, K., Kiyonari, H., and Aizawa, S. (2016). Conserved and divergent expression patterns of markers of axial development in eutherian mammals. Dev Dyn 245, 67–86. 10.1002/dvdy.24352.

47. Idkowiak, J., Weisheit, G., Plitzner, J., and Viebahn, C. (2004). Hypoblast controls mesoderm generation and axial patterning in the gastrulating rabbit embryo. Dev Genes Evol 214, 591–605. 10.1007/s00427-004-0436-y.

48. Latos, P.A., and Hemberger, M. (2016). From the stem of the placental tree: trophoblast stem cells and their progeny. Development 143, 3650–3660. 10.1242/dev.133462.

49. Lau, K.Y.C., Rubinstein, H., Gantner, C.W., Hadas, R., Amadei, G., Stelzer, Y., and Zernicka-Goetz, M. (2022). Mouse embryo model derived exclusively from embryonic stem cells undergoes neurulation and heart development. Cell Stem Cell 29, 1445–1458 e1448. 10.1016/j.stem.2022.08.013.

50. Ben-Haim, N., Lu, C., Guzman-Ayala, M., Pescatore, L., Mesnard, D., Bischofberger, M., Naef, F., Robertson, E.J., and Constam, D.B. (2006). The nodal precursor acting via activin receptors induces mesoderm by maintaining a source of its convertases and BMP4. Dev Cell 11, 313–323. 10.1016/j.devcel.2006.07.005.

51. Donnison, M., Beaton, A., Davey, H.W., Broadhurst, R., L’Huillier, P., and Pfeffer, P.L. (2005). Loss of the extraembryonic ectoderm in Elf5 mutants leads to defects in embryonic patterning. Development 132, 2299–2308. 10.1242/dev.01819.

52. Georgiades, P., and Rossant, J. (2006). Ets2 is necessary in trophoblast for normal embryonic anteroposterior axis development. Development 133, 1059–1068. 10.1242/dev.02277.

53. Navarrete Santos, A., Ramin, N., Tonack, S., and Fischer, B. (2008). Cell lineage-specific signaling of insulin and insulin-like growth factor I in rabbit blastocysts. Endocrinology 149, 515–524. 10.1210/en.2007-0821.

54. Thieme, R., Ramin, N., Fischer, S., Puschel, B., Fischer, B., and Santos, A.N. (2012). Gastrulation in rabbit blastocysts depends on insulin and insulin-like-growth-factor 1. Mol Cell Endocrinol 348, 112–119. 10.1016/j.mce.2011.07.044.

55. Hopf, C., Viebahn, C., and Puschel, B. (2011). BMP signals and the transcriptional repressor BLIMP1 during germline segregation in the mammalian embryo. Dev Genes Evol 221, 209–223. 10.1007/s00427-011-0373-5.

56. Schroder, S.S., Tsikolia, N., Weizbauer, A., Hue, I., and Viebahn, C. (2016). Paraxial Nodal Expression Reveals a Novel Conserved Structure of the Left-Right Organizer in Four Mammalian Species. Cells Tissues Organs 201, 77–87. 10.1159/000440951.

57. Chen, D., Sun, N., Hou, L., Kim, R., Faith, J., Aslanyan, M., Tao, Y., Zheng, Y., Fu, J., Liu, W., et al. (2019). Human Primordial Germ Cells Are Specified from Lineage-Primed Progenitors. Cell Rep 29, 4568–4582 e4565. 10.1016/j.celrep.2019.11.083.

58. Kobayashi, T., Zhang, H., Tang, W.W.C., Irie, N., Withey, S., Klisch, D., Sybirna, A., Dietmann, S., Contreras, D.A., Webb, R., et al. (2017). Principles of early human development and germ cell program from conserved model systems. Nature 546, 416–420. 10.1038/nature22812.

59. Michael, A.K., Harvey, S.L., Sammons, P.J., Anderson, A.P., Kopalle, H.M., Banham, A.H., and Partch, C.L. (2015). Cancer/Testis Antigen PASD1 Silences the Circadian Clock. Mol Cell 58, 743–754. 10.1016/j.molcel.2015.03.031.

60. Levin, M., Anavy, L., Cole, A.G., Winter, E., Mostov, N., Khair, S., Senderovich, N., Kovalev, E., Silver, D.H., Feder, M., et al. (2016). The mid-developmental transition and the evolution of animal body plans. Nature 531, 637–641. 10.1038/nature16994.

61. Puschel, B., and Viebahn, C. (2010). Rabbit mating and embryo isolation. Cold Spring Harb Protoc 2010, pdb prot5350. 10.1101/pdb.prot5350.

62. Viebahn, C., Stortz, C., Mitchell, S.A., and Blum, M. (2002). Low proliferative and high migratory activity in the area of Brachyury expressing mesoderm progenitor cells in the gastrulating rabbit embryo. Development 129, 2355–2365. 10.1242/dev.129.10.2355.

63. Keren-Shaul, H., Kenigsberg, E., Jaitin, D.A., David, E., Paul, F., Tanay, A., and Amit, I. (2019). MARS-seq2.0: an experimental and analytical pipeline for indexed sorting combined with single-cell RNA sequencing. Nat Protoc 14, 1841–1862. 10.1038/s41596-019-0164-4.

64. Dobin, A., Davis, C.A., Schlesinger, F., Drenkow, J., Zaleski, C., Jha, S., Batut, P., Chaisson, M., and Gingeras, T.R. (2013). STAR: ultrafast universal RNA-seq aligner. Bioinformatics 29, 15–21. 10.1093/bioinformatics/bts635.

