## Supplemental figures for "Time-Aligned Hourglass Gastrulation Models in Rabbit and Mouse"

Figure S1

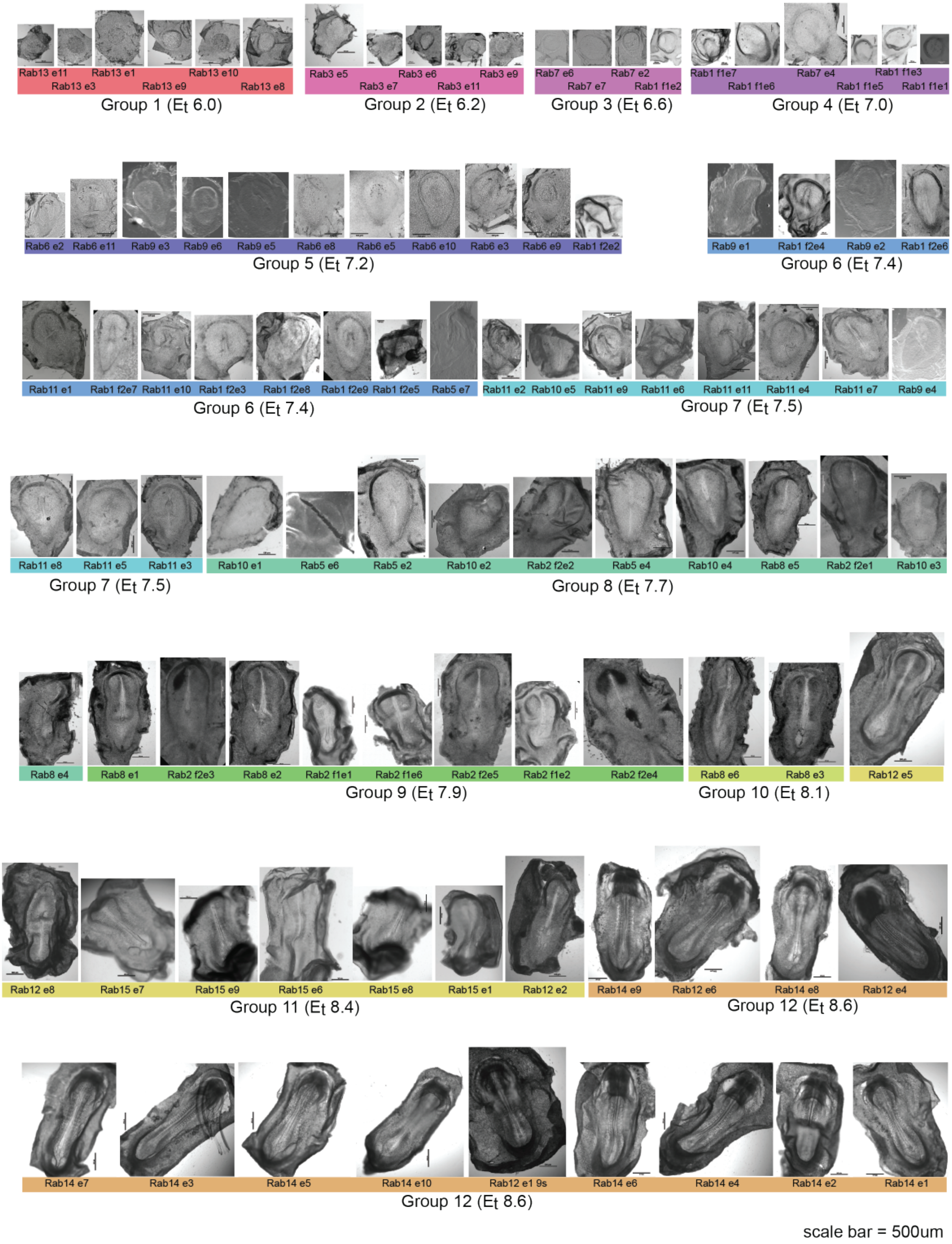
Rab13 e11 Rab13 e1 Rab13 e10
Rab13 e3 Rab13 e9 Rab13 e8
Group 1 (Et 6.0)
Rab3 e5 Rab3 e6 Rab3 e9
Rab3 e7 Rab3 e11
Group 2 (Et 6.2)
Rab7 e6 Rab7 e2
Rab7 e7 Rab1 f1e2
Group 3 (Et 6.6)
Rab1 f1e7 Rab7 e4 Rab1 f1e3
Rab1 f1e6 Rab1 f1e5 Rab1 f1e1
Group 4 (Et 7.0)
Rab6 e2 Rab6 e11 Rab9 e3 Rab9 e6 Rab9 e5 Rab6 e8 Rab6 e5 Rab6 e10 Rab6 e3 Rab6 e9 Rab1 f2e2
Group 5 (Et 7.2)
Rab9 e1 Rab1 f2e4 Rab9 e2 Rab1 f2e6
Group 6 (Et 7.4)
Rab11 e1 Rab1 f2e7 Rab11 e10 Rab1 f2e3 Rab1 f2e8 Rab1 f2e9 Rab1 f2e5 Rab5 e7
Group 6 (Et 7.4)
Rab11 e2 Rab10 e5 Rab11 e9 Rab11 e6 Rab11 e11 Rab11 e4 Rab11 e7 Rab9 e4
Group 7 (Et 7.5)
Rab11 e8 Rab11 e5 Rab11 e3
Group 7 (Et 7.5)
Rab10 e1 Rab5 e6 Rab5 e2 Rab10 e2 Rab2 f2e2 Rab5 e4 Rab10 e4 Rab8 e5 Rab2 f2e1 Rab10 e3
Group 8 (Et 7.7)
Rab8 e4 Rab8 e1 Rab2 f2e3 Rab8 e2 Rab2 f1e1 Rab2 f1e6 Rab2 f2e5 Rab2 f1e2 Rab2 f2e4
Group 9 (Et 7.9)
Rab8 e6 Rab8 e3 Rab12 e5
Group 10 (Et 8.1)
Rab12 e8 Rab15 e7 Rab15 e9 Rab15 e6 Rab15 e8 Rab15 e1 Rab12 e2
Group 11 (Et 8.4)
Rab14 e9 Rab12 e6 Rab14 e8 Rab12 e4
Group 12 (Et 8.6)
Rab14 e7 Rab14 e3 Rab14 e5 Rab14 e10 Rab12 e1 9s Rab14 e6 Rab14 e4 Rab14 e2 Rab14 e1
Group 12 (Et 8.6)
scale bar = 500um

Figure S2

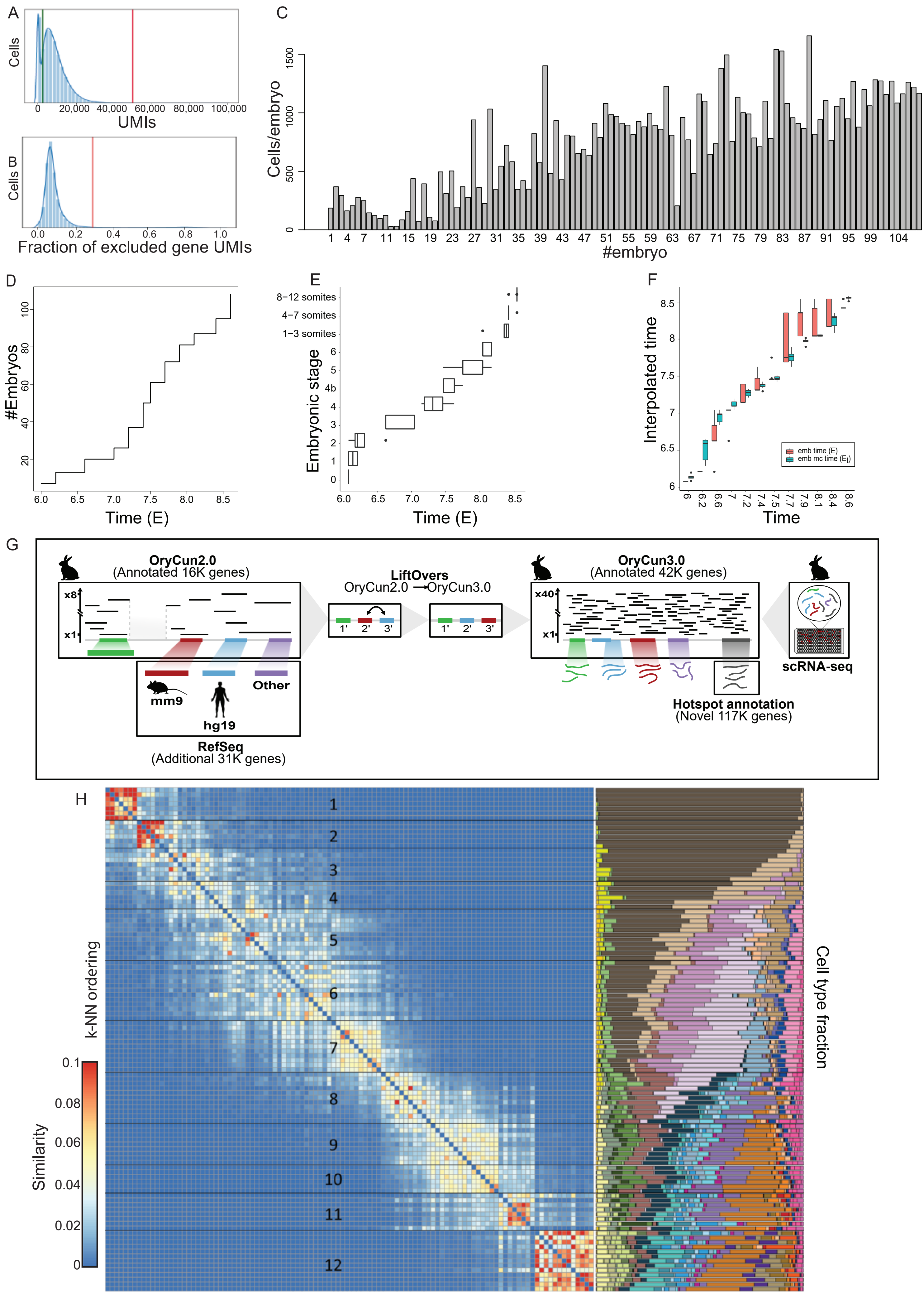
A
Cells
0 20,000 40,000 60,000 80,000 100,000
UMIs
B
Cells
0.0 0.2 0.4 0.6 0.8 1.0
Fraction of excluded gene UMIs
C
Cells/embryo
#embryo
D
#Embryos
Time (E)
E
8−12 somites
4−7 somites
1−3 somites
Embryonic stage
Time (E)
F
Interpolated time
emb time (E)
emb mc time (Et)
Time
G
OryCun2.0
(Annotated 16K genes)
LiftOvers
OryCun2.0 → OryCun3.0
OryCun3.0
(Annotated 42K genes)
x8
x1
x40
mm9
hg19
Other
RefSeq
(Additional 31K genes)
scRNA-seq
Hotspot annotation
(Novel 117K genes)
H
k-NN ordering
Similarity
Cell type fraction

### Figure S3

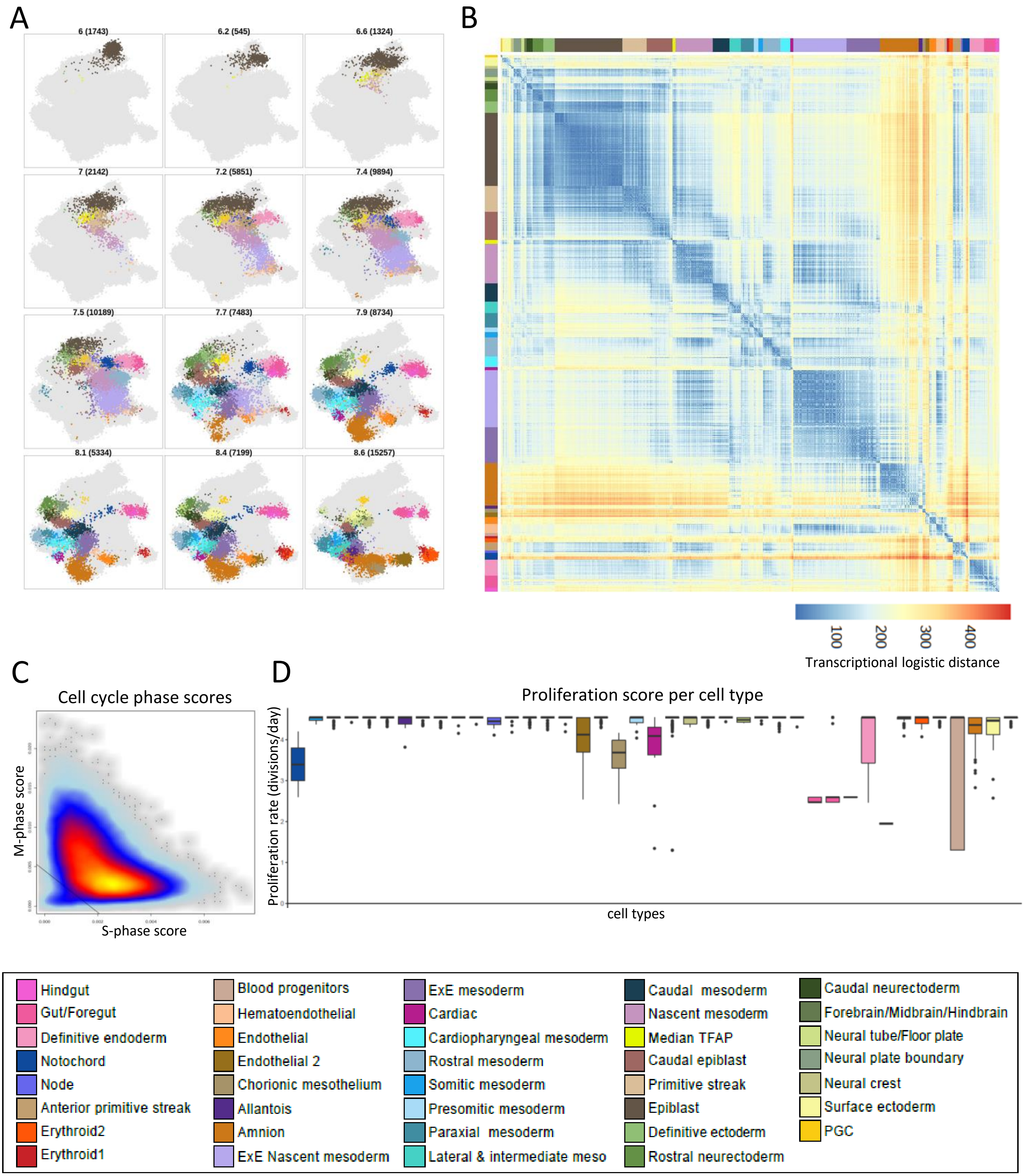
A
6 (1743)
6.2 (545)
6.6 (1324)
7 (2142)
7.2 (5851)
7.4 (9894)
7.5 (10189)
7.7 (7483)
7.9 (8734)
8.1 (5334)
8.4 (7199)
8.6 (15257)
B
100
200
300
400
Transcriptional logistic distance
C
Cell cycle phase scores
M-phase score
S-phase score
D
Proliferation score per cell type
Proliferation rate (divisions/day)
cell types
Hindgut
Gut/Foregut
Definitive endoderm
Notochord
Node
Anterior primitive streak
Erythroid2
Erythroid1
Blood progenitors
Hematoendothelial
Endothelial
Endothelial 2
Chorionic mesothelium
Allantois
Amnion
ExE Nascent mesoderm
ExE mesoderm
Cardiac
Cardiopharyngeal mesoderm
Rostral mesoderm
Somitic mesoderm
Presomitic mesoderm
Paraxial mesoderm
Lateral & intermediate meso
Caudal mesoderm
Nascent mesoderm
Median TFAP
Caudal epiblast
Primitive streak
Epiblast
Definitive ectoderm
Rostral neurectoderm
Caudal neurectoderm
Forebrain/Midbrain/Hindbrain
Neural tube/Floor plate
Neural plate boundary
Neural crest
Surface ectoderm
PGC

### Figure S4

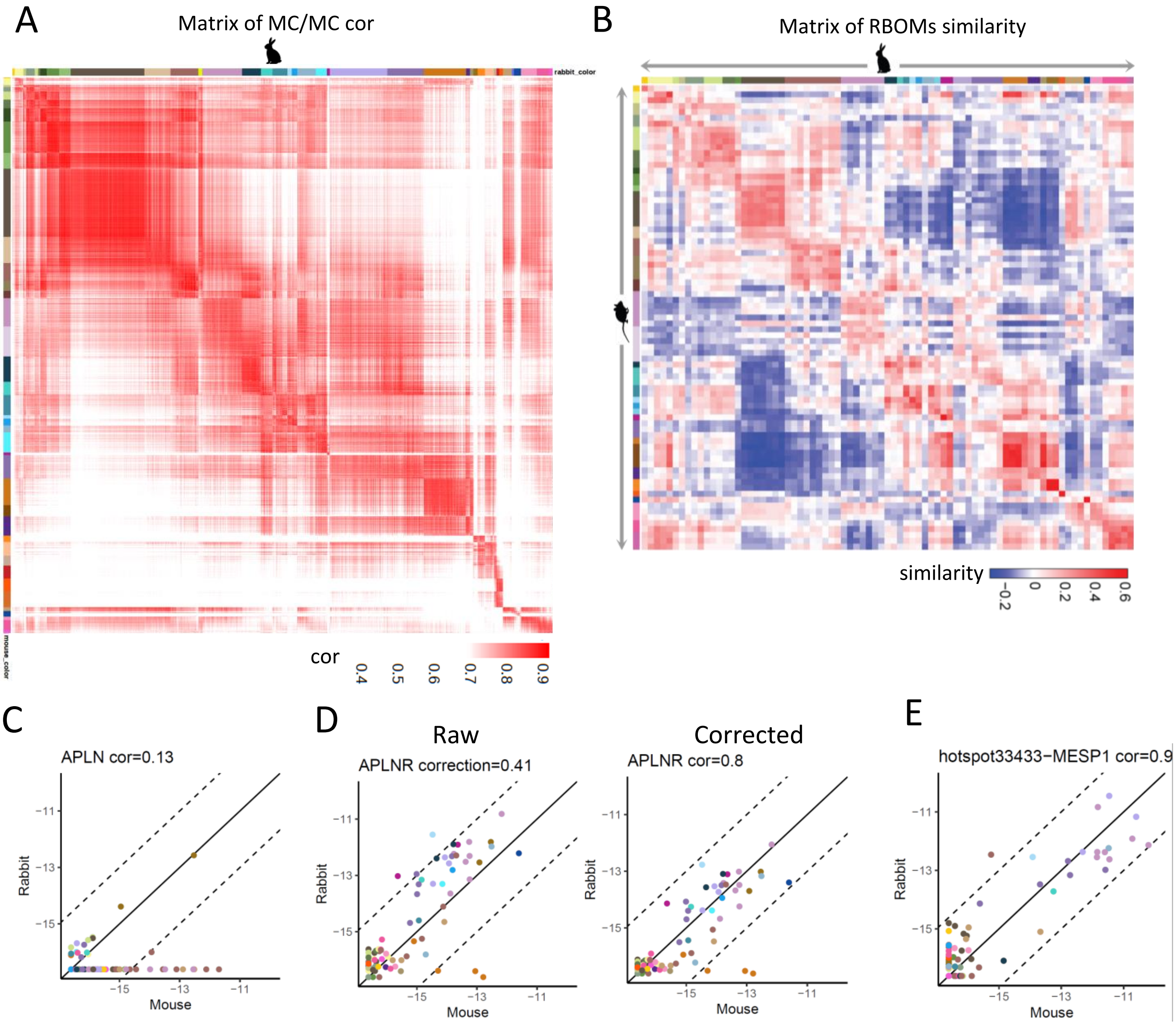


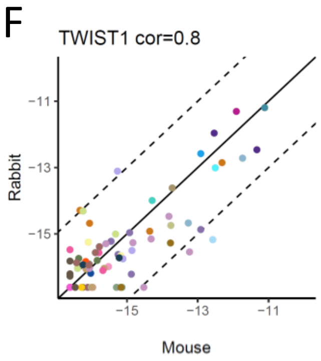


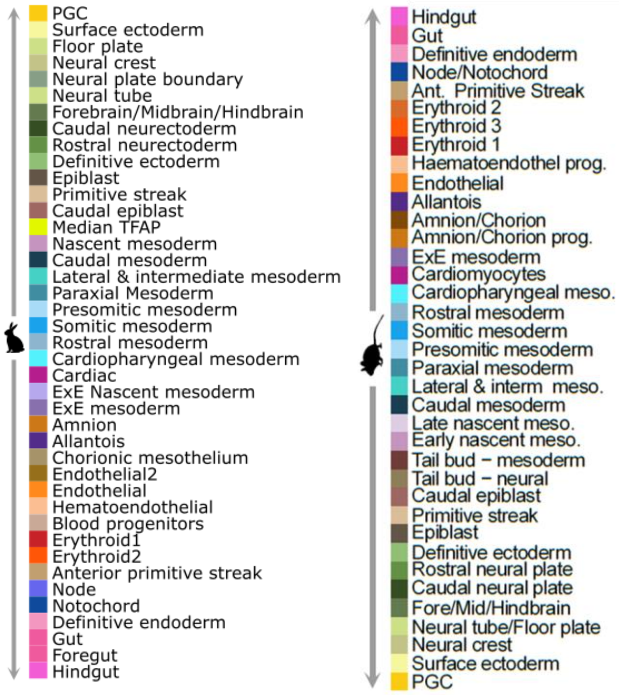


| chromosome | start | end | strand | name |
|---|---|---|---|---|
| chrX | 109106404 | 109109821 | -1 | APLN |
| chrX | 109110464 | 109110636 | -1 | APLN |
| chrX | 109117205 | 109117620 | -1 | APLN |
| chr1 | 6901283 | 6905888 | 1 | APLNR |
| chr17 | 22511500 | 22515900 | 1 | hotspot33433 |
| chr10 | 68567456 | 68567839 | 1 | TWIST1_mm9 |
| chr10 | 68567108 | 68568370 | 1 | TWIST1_mm9 |

Figure S5

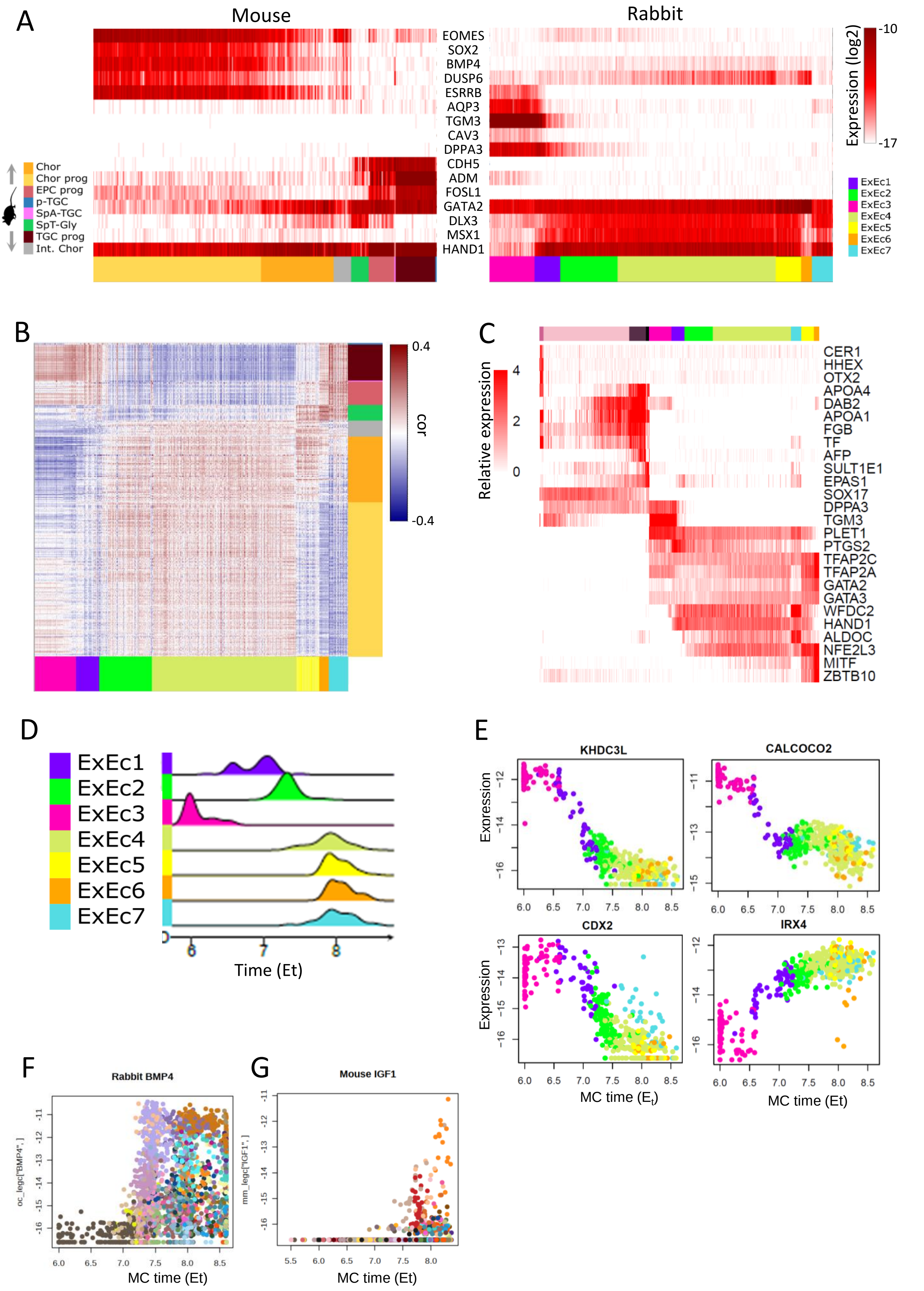

A
Mouse
Rabbit
EOMES
SOX2
BMP4
DUSP6
ESRRB
AQP3
TGM3
CAV3
DPPA3
CDH5
ADM
FOSL1
GATA2
DLX3
MSX1
HAND1
Chor
Chor prog
EPC prog
p-TGC
SpA-TGC
SpT-Gly
TGC prog
Int. Chor
Expression (log2)
-10
-17
ExEc1
ExEc2
ExEc3
ExEc4
ExEc5
ExEc6
ExEc7
B
cor
0.4
-0.4
C
Relative expression
4
2
0
CER1
HHEX
OTX2
APOA4
DAB2
APOA1
FGB
TF
AFP
SULT1E1
EPAS1
SOX17
DPPA3
TGM3
PLET1
PTGS2
TFAP2C
TFAP2A
GATA2
GATA3
WFDC2
HAND1
ALDOC
NFE2L3
MITF
ZBTB10
D
ExEc1
ExEc2
ExEc3
ExEc4
ExEc5
ExEc6
ExEc7
6
7
8
Time (Et)
E
KHDC3L
CALCOCO2
CDX2
IRX4
Expression
MC time (Et)
MC time (Et)
F
Rabbit BMP4
oc_legc["BMP4",]
MC time (Et)
G
Mouse IGF1
mm_legc["IGF1",]
MC time (Et)

Figure S6

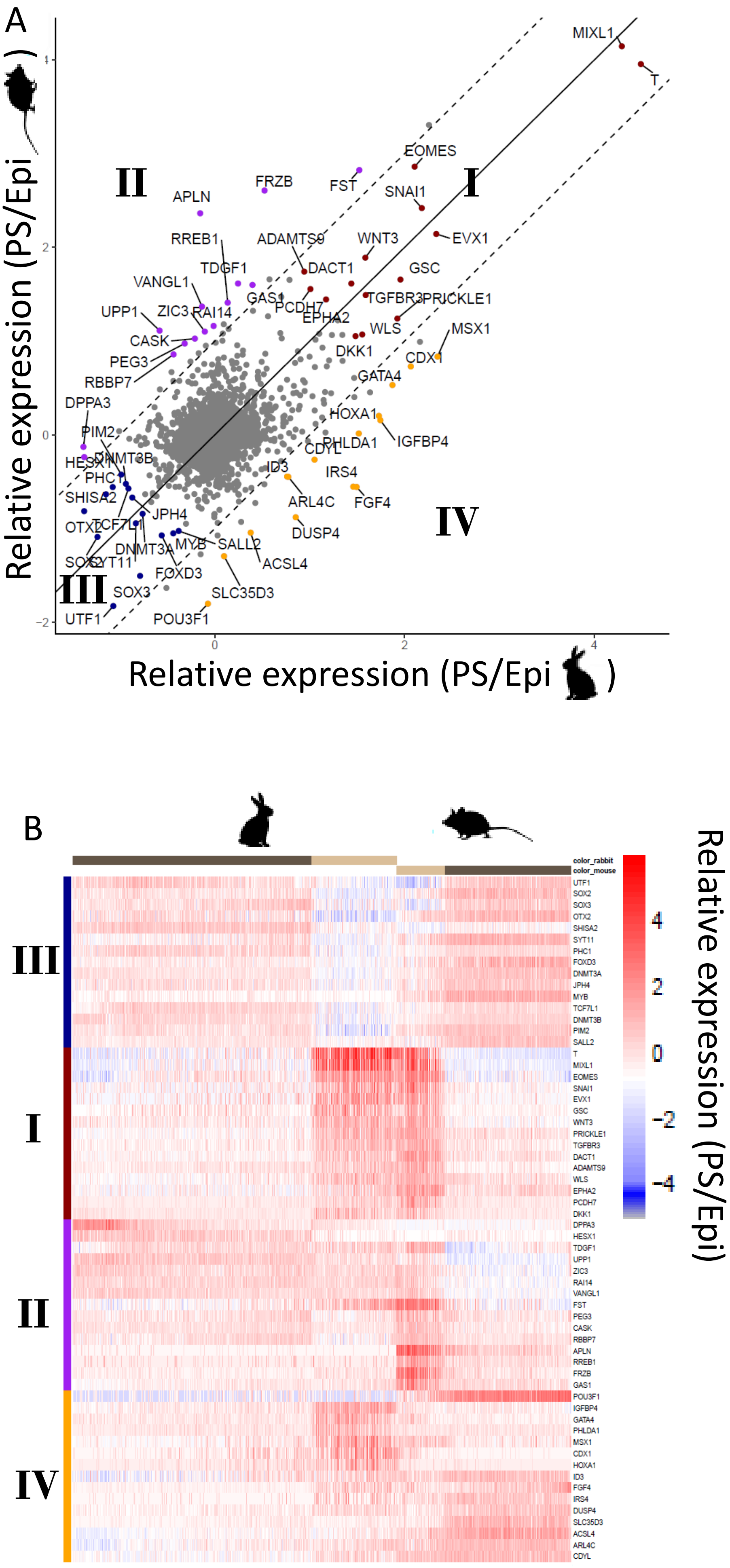

A
B
Relative expression (PS/Epi )
I
II
III
IV
MIXL1
T
EOMES
SNAI1
EVX1
WNT3
GSC
FRZB
FST
APLN
RREB1
ADAMTS9
DACT1
TDGF1
VANGL1
GAS1
PCDH7
EPHA2
TGFBR3
PRICKLE1
UPP1
ZIC3
RAI14
CASK
PEG3
RBBP7
WLS
MSX1
DKK1
CDX1
GATA4
DPPA3
HOXA1
IGFBP4
PIM2
PHLDA1
CDYL
HESX1
DNMT3B
ID3
IRS4
PHC1
SHISA2
ARL4C
FGF4
JPH4
OTX2
TCF7L1
DUSP4
DNMT3A
MYB
SALL2
SOX2
SYT11
ACSL4
FOXD3
SLC35D3
SOX3
UTF1
POU3F1
color_rabbit
color_mouse
Relative expression (PS/Epi)

### Figure S7

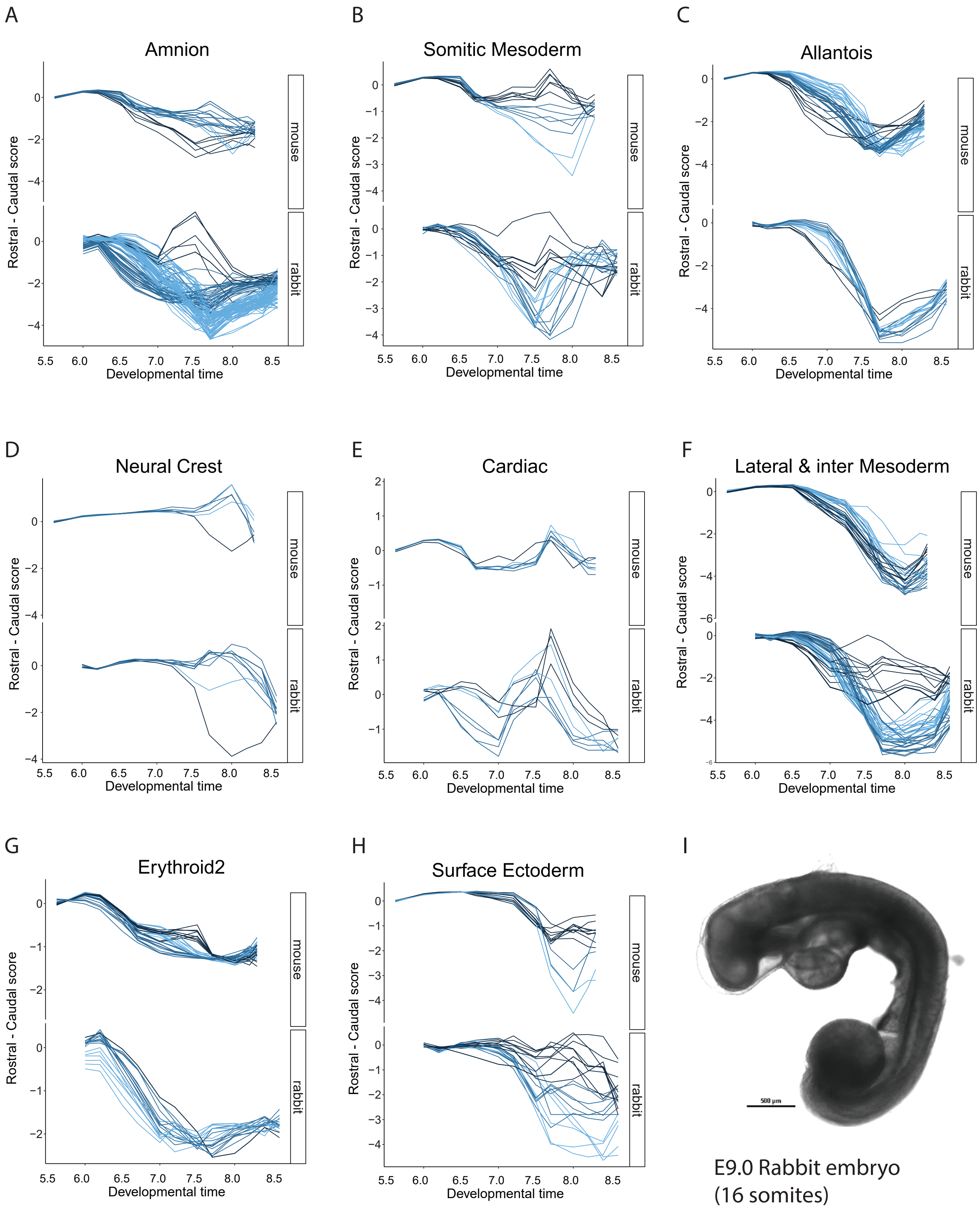

A
Amnion
B
Somitic Mesoderm
C
Allantois
D
Neural Crest
E
Cardiac
F
Lateral & inter Mesoderm
G
Erythroid2
H
Surface Ectoderm
Rostral - Caudal score
Developmental time
5.5
6.0
6.5
7.0
7.5
8.0
8.5
mouse
rabbit
I
E9.0 Rabbit embryo
(16 somites)

Figure S8

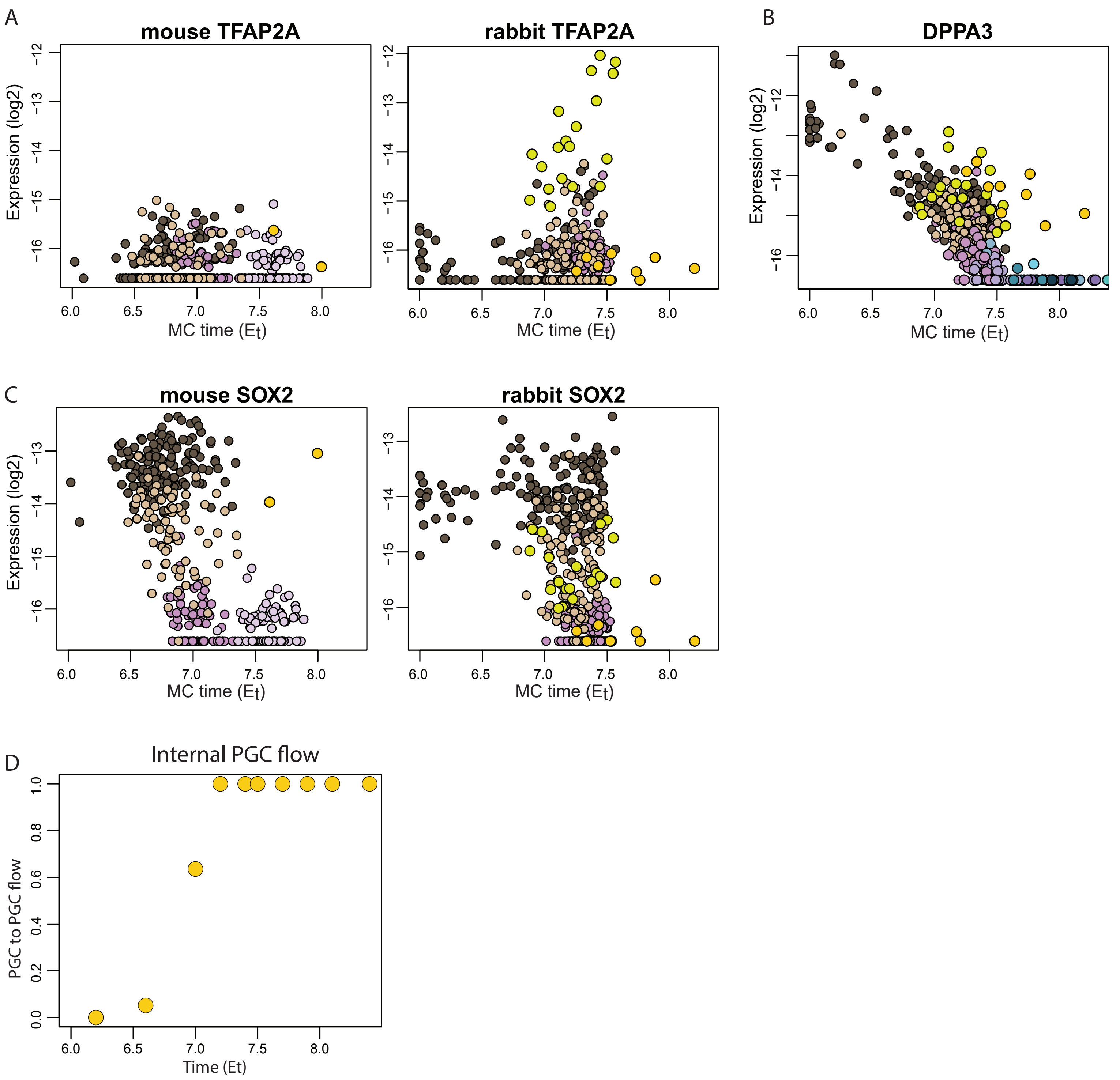
A
mouse TFAP2A
rabbit TFAP2A
B
DPPA3
C
mouse SOX2
rabbit SOX2
D
Internal PGC flow
Expression (log2)
MC time (Et)
PGC to PGC flow
Time (Et)
